# Topologically associating domain (TAD) boundaries stable across diverse cell types are evolutionarily constrained and enriched for heritability

**DOI:** 10.1101/2020.01.10.901967

**Authors:** Evonne McArthur, John A. Capra

## Abstract

Topologically associating domains (TADs) are fundamental units of three-dimensional (3D) nuclear organization. The regions bordering TADs—TAD boundaries—contribute to the regulation of gene expression by restricting interactions of cis-regulatory sequences to their target genes. TAD and TAD boundary disruption have been implicated in rare disease pathogenesis; however, we have a limited framework for integrating TADs and their variation across cell types into the interpretation of common trait-associated variants. Here, we investigate an attribute of 3D genome architecture—the stability of TAD boundaries across cell types—and demonstrate its relevance to understanding how genetic variation in TADs contribute to complex disease. By synthesizing TAD maps across 37 diverse cell types with 41 genome-wide association studies (GWAS), we investigate the differences in functionality and evolutionary pressure on variation in TADs versus TAD boundaries. We quantify their contribution to trait heritability and sequence-level evolutionary constraint and demonstrate that genetic variation in TAD boundaries contributes more to complex trait heritability, especially for immunologic, hematologic, and metabolic traits. We also show that TAD boundaries are more evolutionarily constrained than TADs. Next, stratifying boundaries by their stability across cell types, we find substantial differences. Boundaries stable across cell types are further enriched for complex trait heritability, evolutionary constraint, CTCF binding, and housekeeping genes compared to boundaries unique to a specific cell type. This suggests greater functional importance for stable boundaries. Thus, considering TAD boundary stability across cell types provides valuable context for understanding the genome’s functional landscape and enabling 3D-structure aware variant interpretation.

## INTRODUCTION

The three-dimensional (3D) conformation of the genome facilitates the regulation of gene expression [1–4]. Using chromosome conformation capture technologies (3C, 4C, 5C, Hi-C) [5–7], recent studies have demonstrated that modulation of gene expression via 3D chromatin structure is important for many physiologic and pathologic cellular functions, including cell type identity, cellular differentiation, and risk for multiple rare diseases and cancer [8–13]. Nonetheless, many fundamental questions remain about the functions of and evolutionary constraints on 3D genome architecture. For example, how does genetic variation in different 3D contexts contribute to the risk of common complex disease? Furthermore, disease-causing regulatory variation is known to be tissue-specific; however, only recently has there been characterization of 3D structure variation across multiple cell types and individuals [13–15]. Understanding how different attributes of 3D genome architecture influence disease risk in a cell-type-specific manner is crucial for interpreting human variation and, ultimately, moving from disease associations to an understanding of disease mechanisms [16].

3D genome organization can be characterized at different scales. Globally, chromosomes exist in discrete territories in the cell nucleus [7,17,18]. On a sub-chromosomal scale, chromatin physically compartmentalizes into topologically associating domains (TADs). TADs are megabase-long genomic regions that self-interact, but rarely contact regions outside the domain (Fig. 1A) [7,19–21]. They are likely formed and maintained through interactions between CTCF zinc-finger transcription factors and cohesin ring-shaped complexes, among other proteins both known and unknown [7,22]. TADs are identified as regions of enriched contact density in Hi-C maps (Fig. 1A). TADs modulate gene regulation by limiting interactions of cis-regulatory sequences to target genes [7]. The extent to which chromatin 3D topology affects gene expression is still debated. In extensively rearranged *Drosophila* balancer chromosomes, few genes had expression changes [23]. In contrast, subtle chromatin interaction changes in induced pluripotent stem cells (iPSCs) from seven related individuals were associated with proportionally large differential gene expression [24]. Thus, further cell-type-specific investigation into properties of TAD organization and disruption is needed to clarify which parts of the genome are sensitive to changes in 3D structure.

**Figure 1.**
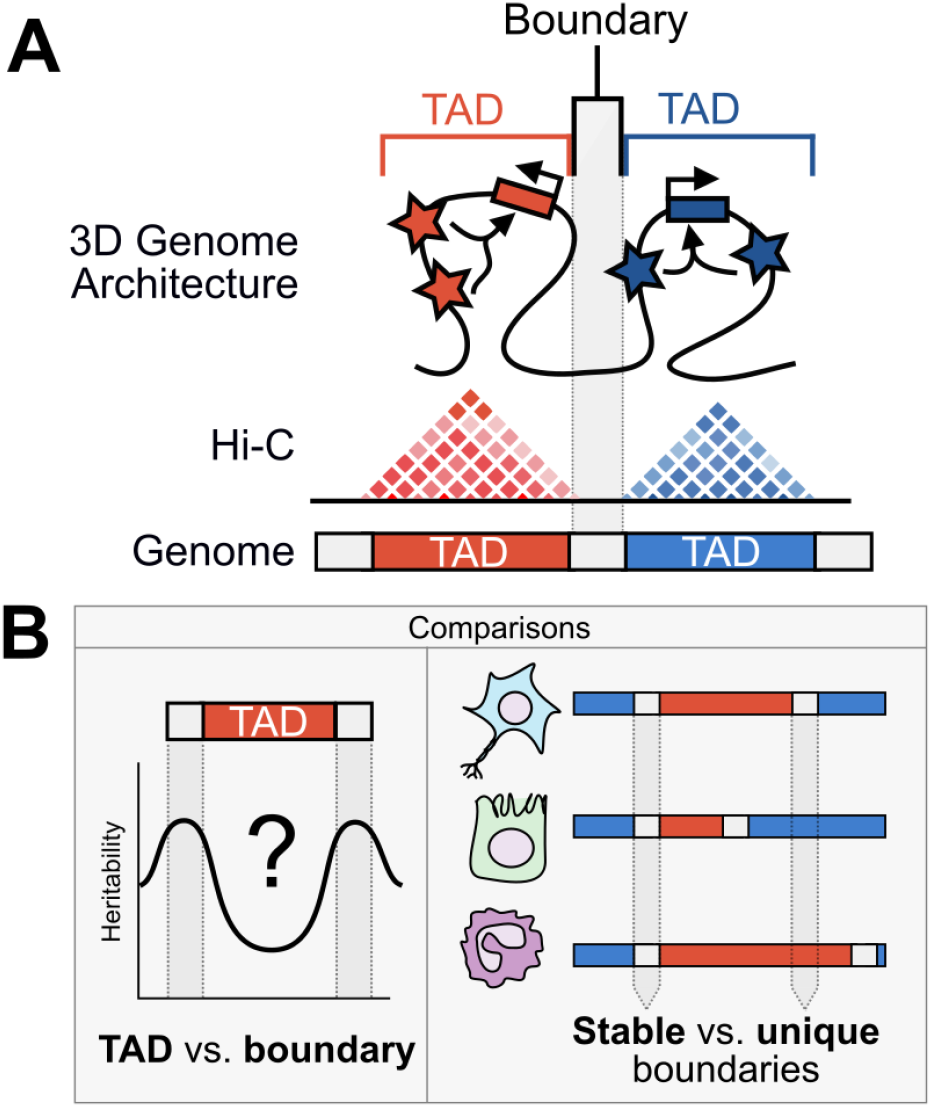
Schematic of our analyses of 3D chromatin TAD boundary stability and functionality. **(A)** Chromatin is organized in 3D space into TADs which are determined by Hi-C experiments. Regions bordering TADs are TAD boundaries. Regions within a TAD are much more likely to interact with one another than regions outside of the TAD. Boxes with right-angled arrows represent genes and stars represent gene regulatory elements, like enhancers. **(B)** This work addresses two main questions: (1) Are TADs or TAD boundaries more enriched for complex trait heritability and evolutionary conservation? (2) Are stable TAD boundaries (*i.e*., those observed across multiple tissues) more or less functionally enriched than TAD boundaries unique to specific tissues?

At the highest level, TAD organization can be divided into two basic features: the TAD and the TAD boundary. TADs are the self-associating, loop-like domains that contain interacting cis-regulatory elements and target genes. TAD boundaries—regions in between TADs—are insulatory elements that restrict interactions of cis-regulatory sequences, like enhancers, to target genes [7]. Previous work suggests the functional importance of maintaining both the “self-associating” TADs and the “insulatory” boundaries. For example, in cross-species multiple sequence alignments, syntenic break enrichment at TAD boundaries suggests a long-term evolutionary preference for rearrangements that “shuffle” intact TADs, rather than “break” them [25,26]. Additionally, TADs often contain clusters of co-regulated genes—*e.g*., cytochrome genes and olfactory receptors [7,20,27]. Intra-TAD structural variation that deletes or duplicates enhancers has been implicated in polydactyly, B cell lymphoma, and aniridia [28]. Together, these data suggest that the genome is under pressure to preserve TADs as functional units.

Other evidence suggests the greater importance of maintaining TAD boundaries. TAD boundaries are enriched for housekeeping genes and transcription start sites [7,19]. Removing insulatory TAD boundaries leads to ectopic gene expression in cultured cells and *in vivo*. For example, TAD structure disruption at the *EPHA4* locus leads to inappropriate rewiring of developmental genes implicated in limb formation defects [7,28,29]. In cancer, large structural alterations that disrupt TAD boundaries cause pathogenic gene expression in acute myeloid leukemia (AML) and medulloblastoma [30,31]. Structural variation (SV) that disrupts TAD boundaries causes gain-of-function, loss-of-function, and misexpression in many forms of rare neurodevelopmental disease [28]. Accordingly, TAD boundaries and CTCF sites have evidence of purifying selection on SV [32,33]. Finally, human haplotype breakpoints do not align with chromatin boundaries, which indicates that recombination may be deleterious at TAD boundaries [34]. Collectively, these suggest that TAD boundaries are functionally important and constrained, especially on the scale of human evolution.

In addition to the need for further characterization of TADs versus TAD boundaries, there is also a gap in our understanding of the variability in TAD organization across cell types. TADs and TAD boundaries have been characterized as largely invariant across cell types [19,20,35–37] and species [7,19,26,38,39]. However, previous pair-wise comparisons of five 3D maps suggest that 30-50% of TADs differ across cell types [36,40]. Boundaries shared across two cell types have evidence of stronger SV purifying selection than “unique” boundaries, suggesting that “stable” boundaries are more intolerant of disruption [32]. Although this provides some characterization of “stable” versus “unique” TAD boundaries across cell types, most previous attempts to categorize TAD boundaries have focused on insulatory strength or hierarchical level in a single cell type. For example, stratifying boundaries by their strength (in a single cell type) facilitated discovery that greater CTCF binding confers stronger insulation and that super-enhancers are preferentially insulated by the strongest boundaries [41]. Stratifying by hierarchical properties of TADs—TADs often have sub-TADs—demonstrated that boundaries flanking higher-level structures are enriched for CTCF, active epigenetic states, and higher gene expression [42].

Despite these preliminary indications that the stability of components of the 3D architecture may influence functional constraint, there has been no comprehensive analysis comparing genomic functionality and disease-associations between 3D structural elements stable across multiple cell types versus those that are unique to single cell types. Quantifying stability across cell types is important for interpreting new variation within the context of the 3D genome given our knowledge that disease-associated regulatory variation is often tissue-specific [13–15].

To investigate differences in TAD boundaries across cell types, we quantify boundary “stability” as the number of tissues that share a TAD boundary. If a TAD boundary is found in many tissues, it is “stable”; whereas, if it is found in few tissues, it is “unique” (Fig. 1B). Using this characterization, we address two main questions that aim to expand our framework for cell-type-aware interpretation of genetic variation and disease associations in the context of the 3D genome (Fig. 1B):

1. How do **TADs** versus **TAD boundaries** differ in terms of their contribution to complex trait heritability and their evolutionary constraint?
2. Are there functional and evolutionary differences in TAD boundaries **stable** across multiple cell types versus TAD boundaries **unique** to specific tissues?

Synthesizing 3D genome maps across 37 diverse cell types with multiple functional annotations and genome-wide association studies (GWAS), we show that TAD boundaries are more enriched for heritability of common complex traits and more evolutionarily conserved compared to TADs. Furthermore, genetic variation in TAD boundaries stable across multiple cell types contributes more to the heritability of immunologic, hematologic, and metabolic traits than variation in TAD boundaries unique to a single cell type. Finally, these cell type stable TAD boundaries are also more evolutionarily constrained and enriched for functional elements. Together, our work suggests that TAD boundary stability across cell types provides valuable context for understanding the genome’s functional landscape and enabling 3D-structural aware variant interpretation.

## RESULTS

### Estimating complex trait heritability across the 3D genome landscape

Disruption of 3D genome architecture has recently been shown to play a role in rare disease and cancer; however, the relationship between 3D genome landscape and common disease has not been investigated. In contrast to Mendelian traits, the heritability of complex polygenic traits is spread throughout the genome, and genetic variation in functional elements, like enhancers, contributes a greater proportion of heritability than variation in less functional regions [43]. Single nucleotide polymorphisms (SNPs) can influence 3D genome structure, for example, by modifying CTCF-binding site motifs necessary for TAD formation [7,22]. However, the contribution of common variation in different 3D contexts to common phenotypes is unknown. We use partitioned heritability analysis to quantify the relationship between different attributes of the 3D genome architecture and the genetic architecture of common phenotypes.

To investigate these heritability patterns across the 3D genome landscape, we use 37 TAD maps from the 3D Genome Browser (Table S1) [44]. The cellular contexts include primary tissues, stem cells, and cancer cell lines; for simplicity, we will refer to these as “cell types” [35–37,45–48]. All TAD maps were systematically predicted from Hi-C data using the hidden Markov model (HMM) pipeline from Dixon *et al*. (2012) at either 40 kb or 25 kb resolution (Supplemental Text) [19,44].

We estimated common trait heritability enrichment among SNPs within these 3D genome annotations using stratified-LD score regression (S-LDSC) [43,49]. S-LDSC is a method of partitioning heritability across the genome using GWAS summary statistics and LD patterns to test whether an annotation of interest (e.g. TADs or TAD boundaries) is enriched for heritability of a trait compared to the rest of the genome. We considered GWAS summary statistics from a previously-described representative set of 41 diseases and complex traits (average N = 329,378, M = 1,155,239, h^2^_SNP_ = 0.19, Table S2) [50,51,60,52–59].

Motivated by the approach to partitioning TADs from Krefting *et al*., we analyzed TADs plus 50% of their total length on each side and subdivided this region into 20 equal-sized partitions [25]. Bins 1-5 and 16-20 “bookend” the TAD, while the center bins 6-15 are inside the TAD. In cases where a TAD is in very close proximity to another TAD, the ±50% region flanking the TAD (bins 1-5,16-20) may extend into a neighboring TAD. However, generally, TADs represent the center 10 partitions and we define TAD boundaries as the partitions that flank the TAD. We then conducted S-LDSC for each of the 20 partitions across the 37 cell types for the 41 traits to estimate the enrichment (or depletion) of heritability for that trait across the TAD landscape.

### TAD boundaries are enriched for complex trait heritability and evolutionary conservation

TAD boundaries are significantly enriched for complex trait heritability; whereas TADs are marginally depleted for heritability overall (1.07x enrichment in boundaries vs. 0.99x enrichment in TADs, *P* ~ 0) (Fig. 2A). There is a spike of heritability enrichment centrally in TADs; we explore this further in a subsequent section. The results are consistent whether averaged across traits or meta-analyzed using a random-effects model [43,51,61] (*r*^2^ = 0.85, *P* = 7×10^−9^, Fig. S1); therefore, further analyses of heritability across traits will use averaging for simplicity and interpretability.

**Figure 2.**
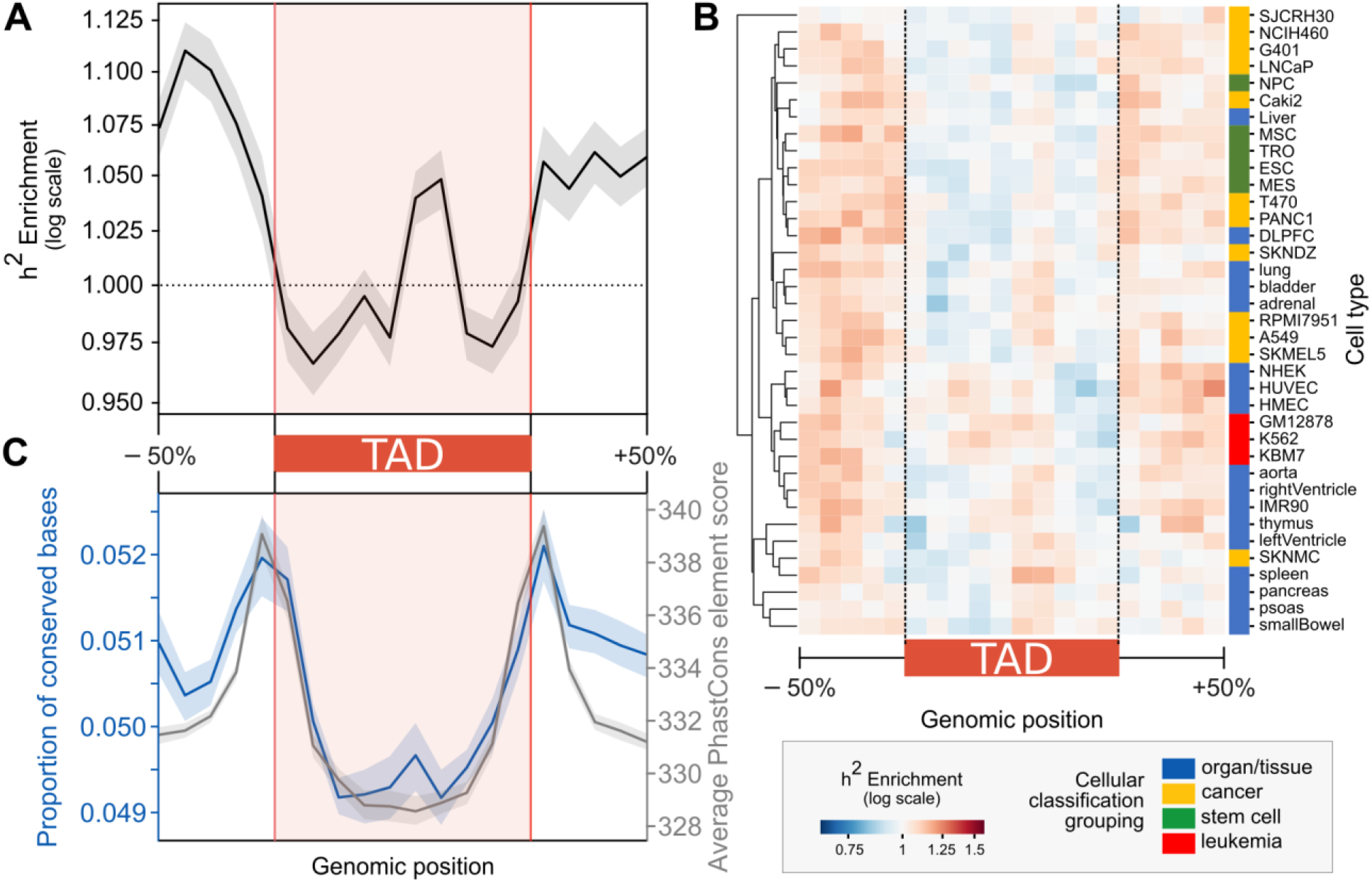
TAD boundaries are enriched for heritability of diverse common complex traits and evolutionary conservation. **(A)** For TADs across 37 cell types, heritability is enriched near TAD boundaries and in the center of TADs when averaged across 41 common complex phenotypes (*P* ~ 0). **(B)** TAD boundary heritability patterns are consistent across the 3D genome landscape for 37 cell types. **(C)** Compared to the interior of TADs, TAD boundaries have increased sequence-level constraint. TAD boundaries have a higher proportion of conserved bases (overlap with PhastCons elements, *P* = 5×10^−55^) (left blue axis) and, across those overlapping PhastCons elements, TAD boundaries have a higher average conservation score (right gray axis) (3×10^−29^). Error bands signify 99% confidence intervals.

The complex trait heritability enrichment at TAD boundaries is also consistent across cell types (Fig. 2B). The heritability enrichment values are highly significant, but as expected, relatively small in magnitude given the large genomic regions considered by this analysis—only a small fraction of the base pairs in a boundary are likely to be functionally relevant.

To assess functionality using a complementary metric, we quantified between-species sequence-level conservation for TADs and boundaries. TAD boundary sequences are significantly more evolutionarily conserved than sequences in TADs (Fig. 2C). We quantified evolutionary conservation in terms of the proportion of base pairs in a region in a conserved element identified by PhastCons elements and by the average PhastCons element score across the region. On average, 5.11% of TAD boundaries are overlapped by PhastCons elements, compared to 4.98% overlapping TADs (*P* = 5×10^−55^). This aligns with previous findings underscoring the importance of maintaining TAD boundaries.

The heritability enrichment and conservation at TAD boundaries are likely due to their overlap with functional elements like CTCF binding sites and genes. Many such elements are enriched for heritability themselves [43]. To assess whether the TAD boundary heritability enrichment is greater than expected given the known functional elements overlapping TAD boundaries, we calculate standardized enrichment effect sizes (τ*) [51,62]. This metric quantifies heritability unique to the focal annotation by conditioning on a broad set of 86 functional regulatory, evolutionary conservation, coding, and LD-based annotations (baseline v2.1) [43,51,62,63]. TAD boundaries did not show more heritability than expected based on their enrichment for the 86 other annotations (Fig. S2). Thus, existing annotations likely capture relevant functional elements (*e.g*., CTCF, conservation, and other regulatory element binding sites) that determine and maintain boundary function.

### TAD boundaries vary in stability across cellular contexts

Although the heritability enrichment patterns are similar across cell types and TADs have been characterized as largely invariant across cell types [19,20,35–37], previous work suggests distinct functional properties among TAD boundaries with different insulatory strengths, hierarchical structures, and cell types of activity [32,41,42]. Thus, we hypothesized that the stability of TAD boundaries across cell types would be informative about their functional importance and conservation. To characterize the stability of TAD boundaries across diverse cellular contexts, we defined TAD boundaries as the 100 kb window bookending the TADs (described above) across 37 cell types. Since the maps for each cell type are defined with respect to the same 100 kb windows across the genome, we identify shared, or “stable”, boundaries based on these 100 kb windows (Fig. 3A). We chose to investigate 100 kb boundaries because this corresponds to the +/-10% windows surrounding the TAD where we observe prominent heritability enrichment (Fig. 2A-B) (the median TAD length is 1.15 Mb [IQR: 0.71 - 1.82 Mb]). However, our results are robust to different definitions of TAD boundaries including a 40 kb window surrounding (± 20 kb) TAD start and stop sites (“40 kb boundaries”) and 200 kb windows flanking the TAD start and stop sites (“200 kb bookend boundaries”) (Fig. S3, Methods).

**Figure 3.**
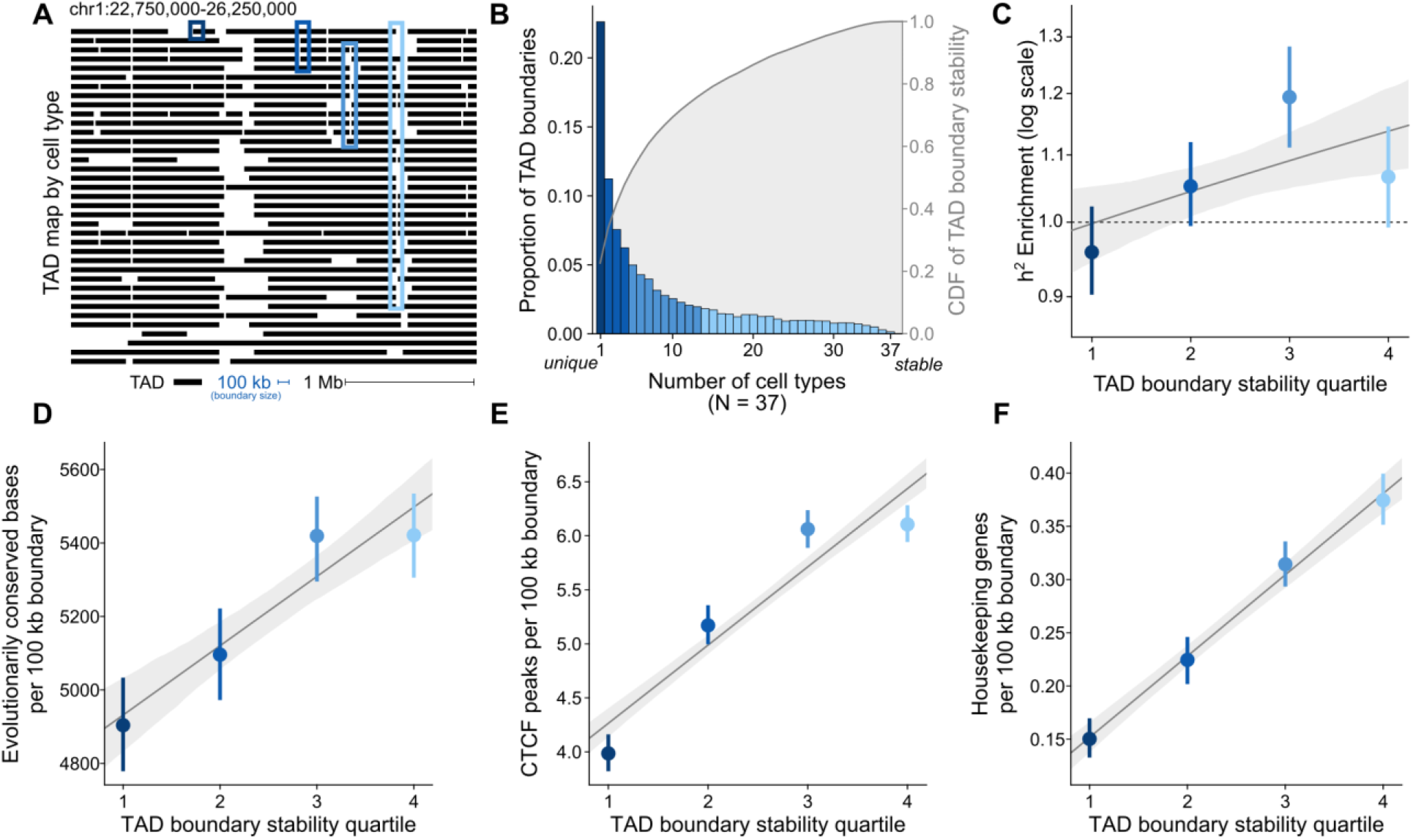
Stable TAD boundaries are enriched for complex trait heritability, evolutionary conservation, and functional elements. **(A)** 37 cell type TAD maps (rows) for a 3.5 Mb window from human chr1 (hg19). Each black line represents the extent of a TAD. Example boundaries of different stability quartiles are outlined in blue (quartile 1 [most cell type unique] in the darkest blue and quartile 4 [most cell type stable] in light blue). **(B)** Histogram of TAD boundaries by the number of cell types they are observed in (their “stability”) colored by quartiles. The right axis and gray distribution represent the empirical cumulative distribution function (CDF) of boundary stability shown in the histogram. Across TAD boundary stability quartiles, there is a correlation between increased cell type stability and increased **(C)** complex trait heritability enrichment (*P* = 0.006), **(D)** conserved bases (overlap with PhastCons elements, *P* = 6×10^−13^), **(E)** CTCF binding (overlap with ChIP-seq peaks, *P* = 1×10^−83^), and **(F)** housekeeping genes (*P* = 8×10^−58^). All error bars/bands signify 95% confidence intervals.

Using the cross-cell-type TAD boundary intersection, we find that boundaries vary substantially across cell types. Less than 10% of TAD boundaries are shared in 25+ of the 37 cell types, and 22.6% of TAD boundaries are unique to a single cell type (Fig. 3B). With the more granular 40 kb boundaries, 33.9% of boundaries are unique to one tissue (Fig. S3A). Even with the permissive 200 kb resolution boundaries, 18.3% of boundaries are unique to a single tissue. (Fig. S3B). To quantify boundary stability for further analyses, we bin boundaries into their cell type stability quartile. For example, boundaries present in only one context of 37 (cell type unique) are in the first quartile of stability, boundaries in 2-4 cell types are in the second quartile, boundaries in 5-13 cell types are in the third quartile, and boundaries in 14 or more of the 37 contexts are the fourth quartile of cell type stability (Fig. 3B, examples in Fig. 3A).

Although there is high variability in TAD boundary landscape across different cell types, we found that biologically similar cell types have more similar TAD boundary maps. For example, cell type classes (*e.g*. organ/tissue, stem cell, and cancer) generally cluster together. The two neuroblastoma cell lines cluster together, as do left ventricle, right ventricle, aorta, and skeletal muscle (Fig. S4B). This trend of biologically similar clusters held at both the 40 kb and 200 kb boundary resolution (Figs. S4A,C). Previous studies have found contrasting results about the level and patterns of similarity across cell types (Supplemental Text), but our similarity quantifications between cell types agree with previous estimates.

In summary, although TADs and TAD boundaries are characterized as largely invariant across cell types, we demonstrate that there is substantial 3D variability between cell types [19,20,35–37]. We also find that biologically related cell types have more similar TAD maps, providing preliminary evidence for the cell type specificity of the 3D genome and providing further rationale for investigating TAD map cell type differences.

### Stable TAD boundaries are enriched for complex trait heritability, evolutionary constraint, and functional elements

When stratifying the 100 kb boundaries by their cell type stability we find a modest, but significant positive relationship between cell-type-stability and trait heritability enrichment (*r*^2^ = 0.045, *P* = 0.006, Fig. 3C). The most stable boundaries (fourth quartile, darkest blue) have 1.07x enrichment of trait heritability compared to 0.96x enrichment in unique boundaries (first quartile). This positive relationship between heritability and boundary stability holds at both the 40 kb and 200 kb resolution (Fig. S5A,D).

We also explored the relationship between TAD boundary stability and other evolutionary and functional attributes. Although TAD boundaries, when compared to TADs, are enriched for CTCF binding [19,41], metrics of evolutionary constraint (Fig. 2C, [32,34]), and housekeeping genes are enriched at TAD boundaries [7,19] (compared to TADs), it is unknown whether these features play a role in boundary stability across cell types.

We find that TAD boundary stability is positively correlated with increased sequence level constraint (Fig. 3D, *P* = 6×10^−13^); boundaries in the highest quartile of stability have an additional 517 base pairs overlap with PhastCons elements compared to cell type unique TAD boundaries (5421 vs 4904 per 100 kb boundary). This extends previous observations that investigated two cell types to show that shared have evidence of stronger purifying selection on structural variants than “unique” boundaries [32]. Based on this, we argue that “stable” boundaries are more intolerant of disruption, not only on the scale of structural variants, but also at the base pair level.

TAD boundary stability is also correlated with increased CTCF binding (Fig. 3E, *P* = 1×10^−83^); for example, boundaries in the highest quartile of stability have 1.5x more CTCF sites on average compared to TAD boundaries unique to one cell type (6.1 vs 4.0). CTCF binding was determined through CTCF ChIP-seq analyses obtained from ENCODE [47,48]. This aligns with previous findings that boundary insulatory strength (in a single cell type) is positively associated with CTCF binding [19,41]; however, it expands this finding to stability across cell types.

Finally, we find that TAD boundary stability is correlated with increased overlap with genes (Fig. S6A-C, *P* = 1×10^−74^), protein-coding genes (Fig. S6D-F, *P* = 7×10^−90^), and housekeeping genes (Figs. 3F, S6G-I, *P* = 8×10^−58^). Boundaries in the highest quartile of stability overlap 2.5x more housekeeping genes compared to cell type unique TAD boundaries (0.37 vs 0.15 per 100 kb boundary). The relationship between stable TAD boundaries and housekeeping gene enrichment may result from many factors, including strong enhancer-promoter interactions, specific transcription factor binding, or because highly active sites of transcription may cause chromatin insulation [12].

In summary, TAD boundaries stable across multiple cell types are enriched for complex trait heritability, evolutionary constraint, CTCF binding, and housekeeping genes. These trends hold at different boundary definitions (40 kb and 200 kb) and for other metrics of conservation, CTCF binding, and gene overlap (Figs. S6-S8).

### The heritability landscape across the 3D genome varies across phenotypes

The previous analyses have shown that trait heritability is generally enriched at TAD boundaries and further enriched in boundaries stable across cell types. Given preliminary evidence that different traits have unique enrichment profiles among different functional annotations [43], we hypothesized that variation in TAD boundaries may influence certain traits more than others. To investigate trait-specific heritability across the TAD landscape, we computed heritability enrichment profiles across the 3D genome partitions by trait and hierarchically clustered them (Fig. 4A). We observed two distinct trait clusters (Fig. 4A).

**Figure 4.**
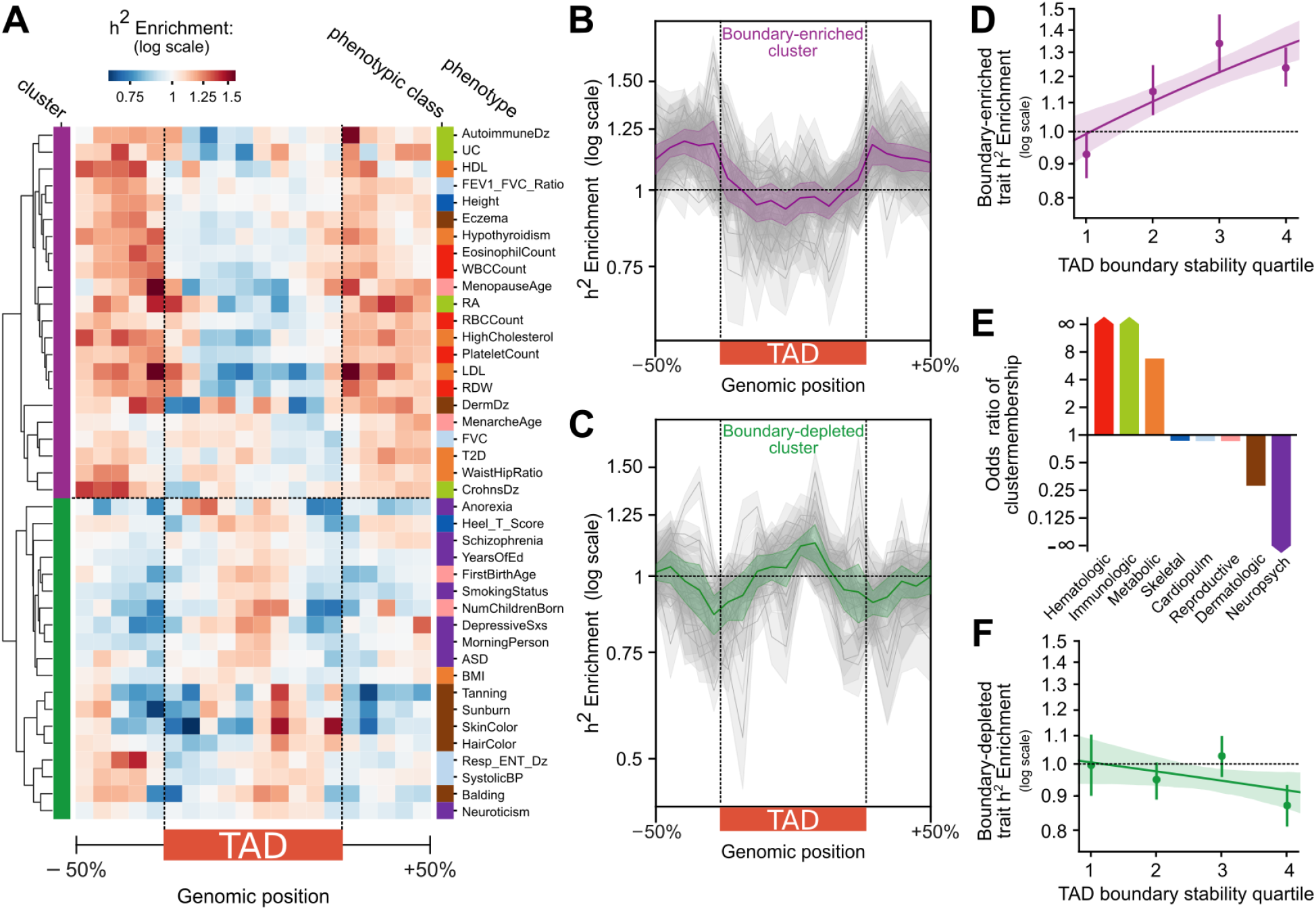
The heritability landscape across the 3D genome varies across phenotypes. **(A)** TAD boundary trait heritability patterns across the 3D genome organize into two clusters. Some traits are strongly enriched for complex trait heritability at TAD boundaries (“boundary-enriched” cluster, purple), while others are weakly depleted at TAD boundaries and enriched centrally within the TAD (“boundary-depleted” cluster, green). **(B)** Heritability enrichment landscape over TADs for traits in the boundary-enriched cluster (N = 22). The gray lines represent the heritability pattern for each trait in the cluster; the purple line is the average over all the traits. **(C)** Heritability enrichment landscape over TADs for traits in the boundary-depleted cluster (N = 19). The gray lines represent the heritability pattern for each trait in the cluster; the green line is the average over all the traits. **(D)** The positive correlation between boundary stability and trait heritability (Fig. 3C) is even stronger for the subset of traits in the boundary-enriched cluster (*r*^2^ = 0.23, *P* = 2×10^−6^). **(E)** Odds of cluster membership across phenotype categories. The boundary-enriched cluster is predominantly hematologic, immunologic, and metabolic traits. The boundary-depleted cluster is predominantly neuropsychiatric traits. **(F)** There is a weak negative correlation between boundary stability and trait heritability for traits in the boundary-depleted cluster (*r*^2^ = 0.04, *P* = 0.09). Error bars/bands signify 99% confidence intervals in B & C, 95% confidence intervals in D & F.

One cluster of traits (“boundary-enriched” cluster) is strongly enriched for complex trait heritability at TAD boundaries (Fig. 4B). Across TAD maps in 37 cell types, these traits have on average 1.15x heritability enrichment at TAD boundaries compared to genomic background and 1.18x enrichment compared to TADs (*P* ~ 0). The other cluster of traits (“boundary-depleted” cluster) shows a weak inverted pattern compared to the boundary-enriched cluster, with marginal heritability depletion at TAD boundaries (0.96x enrichment, *P* ~ 0) and a spike of heritability enrichment within the TAD center (Fig. 4C).

The traits in the boundary-enriched cluster are predominantly hematologic (*e.g*. white blood cell and red blood cell counts), immunologic (*e.g*. rheumatoid arthritis, Crohn’s disease), and metabolic traits (*e.g*. type 2 diabetes, lipid counts) (Fig. 4E). The traits in the boundary-depleted cluster are mostly neuropsychiatric (*e.g*. schizophrenia, years of education, Autism spectrum disorder) and dermatologic (*e.g*. skin color, balding) (Fig. 4E). Our stratification of complex diseases into phenotypic classes does not perfectly reflect the traits’ pathophysiology. For example, some dermatologic traits fall into the boundary-enriched cluster. However, these dermatologic traits, like eczema, also have a significant immunologic and hematologic basis, which are hallmarks of other traits in the boundary-enriched cluster. Additionally, body mass index (BMI) clustered with the psychiatric-predominant boundary-depleted cluster instead of with other metabolic traits in the boundary-enriched cluster. This is interesting given previous findings that BMI heritability is enriched in central nervous system (CNS)-specific annotations rather than metabolic tissue (liver, adrenal, pancreas) annotations [43]. Skeletal, cardiopulmonary, and reproductive traits do not consistently segregate into one of the clusters (Fig. 4E). This is likely because of the small sample size and heterogeneity of traits in these phenotypic classes.

The relationship between heritability enrichment in TAD boundaries and the trait clusters is not confounded by GWAS trait sample size (N), number of SNPs (M), or the traits’ SNP-based heritability (h^2^_SNP_) (Fig. S9). Despite using a diverse set of cell types, we recognize that the heritability pattern differences between traits could be affected by the representation of cell types investigated. However, given that the pattern of heritability enrichment is consistent across all cell types (Fig. 2B), we are confident that no single cluster of cell types is driving the heritability pattern differences between traits. Furthermore, these patterns are maintained even when calling TADs using a variety of computational methods (Armatus, Arrowhead, DomainCaller, HiCseg, TADbit, TADtree, TopDom), suggesting that the finding of immunologic and hematologic heritability enrichment at TAD boundaries is robust to technical variation (Fig. S10).

Although analysis across all traits revealed a positive relationship between boundary cell-type-stability heritability enrichment (Fig. 3C), we found that this trend differs between the two trait clusters. Traits in the boundary-enriched cluster have further heritability enrichment in cell-type-stable boundaries (*r*^2^ = 0.23, *P* = 2×10^−6^, Fig. 4D). The most stable boundaries (fourth quartile) have 1.23x enrichment of trait heritability compared to 0.93x enrichment in unique boundaries (first quartile). In contrast, traits in the boundary-depleted cluster have a non-significant negative relationship between stability and heritability (*r*^2^ = 0.04, *P* = 0.09, Fig. 4F).

## DISCUSSION

Although we are beginning to understand the role of 3D genome disruption in rare disease and cancer, we have a limited framework for integrating maps of 3D genome structure into the study of genome evolution and the interpretation of common disease-associated variation. Here, we show that TAD boundaries are enriched for common complex trait heritability compared to TADs. Furthermore, in exploring TAD boundaries stable across cell types, we find they are further enriched for heritability of hematologic, immunologic, and metabolic traits, as well as evolutionary constraint, CTCF binding, and housekeeping genes. These findings demonstrate a relationship between 3D genome structure and the genetic architecture of common complex disease and reveal differences in the evolutionary pressures acting on different components of the 3D genome.

Previous work has predominantly characterized the importance and evolutionary constraint of different components of the 3D genome from the perspective of SV and rearrangement events. We address functionality at the level of common single nucleotide variants and evolutionary constraint within humans (~100,000 ya) and across diverse vertebrate species (~13-450 mya).

At the scale of common human variation, we show that TAD boundaries are enriched for SNPs that account for the heritability of common complex traits. By demonstrating this relationship between 3D genome structure and common disease-associated variation, we suggest that TAD boundaries are more intolerant of variation than TADs. This aligns with the finding of Whalen et al. (2019) that human haplotype breakpoints—which are associated with increased variation due to the mutagenic properties of recombination—are depleted at chromatin boundaries [34].

Over vertebrate evolution, we show that TAD boundaries have more sequence-level constraint than TADs. This provides a complementary perspective to Krefting *et al*. (2018) which found that TAD boundaries are enriched for syntenic breaks between humans and 12 vertebrate species, thus, concluding that intact TADs are shuffled over evolutionary time [25]. While shuffling a TAD may “move” its genomic location, preserving the TAD unit also requires maintaining at least part of its boundary. Our work suggests that even though TADs are shuffled, the boundary-defining sequences are under more constraint than the sequences within the TAD. This is further supported by the high TAD boundary concordance between syntenic blocks in different species and depletion of SVs at TAD boundaries in humans and primates [7,19,26,32,38,39].

Slight variation in 3D structure can cause proportionally large changes in gene expression [24]. For example, CTCF helps maintain and form TAD boundaries; consequently, altering CTCF binding often leads to functional gene expression changes, *e.g*., oncogenic gene expression in gliomas [27]. We hypothesize that altering gene regulation though common variant-disruption of transcription-factor motifs important in 3D structure organization, like CTCF, contributes to the enrichment for complex disease heritability. However, variation at TAD boundaries may also disrupt regulatory elements like enhancers, that are known to be enriched at boundaries, without disrupting the TAD architecture. A deeper mechanistic understanding of TAD formation will be critical to further understanding how TAD boundary disruption contributes to both rare and common disease at potentially nucleotide-level and cell type resolution.

Our finding of divergent patterns of TAD boundary heritability enrichment for different traits (enrichment for hematologic, immunologic, and metabolic traits versus depletion for psychiatric and dermatologic) suggests that the 3D genome architecture may play differing roles in the genetic architecture of different traits. As a preliminary test of this hypothesis, we evaluated the relationship between boundary stability and intra-TAD heritability enrichment. We find that, for traits with heritability depletion at boundaries (psychiatric, dermatologic traits), increasingly stable TAD boundaries insulate increased intra-TAD heritability (Fig. S11). For these traits, we speculate that stable boundaries may function to insulate important intra-TAD elements (*e.g*. enhancers or genes). This idea is consistent with previous work showing that super-enhancers are insulated by the strongest boundaries (in a single cell type) [41]. However, for the boundary-enriched traits (hematologic, immunologic, metabolic), we hypothesize that functional elements are located at the stable boundaries (rather than inside the TAD). This is supported by previous work that detected a positive association between genome-wide binding of CTCF, a transcription factor intimately involved in TAD boundary formation, and eczema, an immunologic trait that we identified as part of the boundary-enriched trait cluster [64]. Thus, it will be important to further explore how TAD boundaries (or other functional elements at TAD boundaries) may play different regulatory roles in different traits and diseases.

Finally, we identify substantial variation among 3D maps across cell types. While TAD stability across cell types is greater than expected by chance, our findings expand the number and diversity of cell types compared and identify a large proportion of boundaries unique to single cell types (see Supplemental Text). Furthermore, using our metric of cell-type-stability to stratify TAD boundaries identifies meaningful biological differences: stable boundaries are enriched for common trait heritability, evolutionary constraint, and functional elements.

Our analysis is, however, limited by Hi-C data availability. Despite analyzing TADs identified by the same pipeline, there could be batch or protocol-specific effects because the Hi-C data were generated by different groups. Our results are also contingent on the methods we used to define TADs and the particular cell types considered. This underscores the need for higher resolution Hi-C across replicates of diverse cell types and continued efforts to integrate data from multiple TAD-calling algorithms to more precisely define TAD boundaries, especially given their hierarchical nature [42,65]. Despite these complexities in identifying TAD boundaries, our findings replicate with all our boundary definitions and with different TAD calling pipelines considered.

## CONCLUSIONS

Here, we introduce a metric for the stability of a TAD boundary across cell types and demonstrate enrichment of complex trait heritability, sequence-level constraint, and CTCF binding among stable TAD boundaries. Our work suggests the utility of incorporating 3D structural data across multiple cell types to aid context-specific non-coding variant interpretation. Starting from this foundation, much further work is needed to elucidate the molecular mechanisms, evolutionary history, and cell-type-specificity of TAD structure disruption. Furthermore, while we have identified properties of TAD boundaries stable across cell types, it would also be valuable to identify differences in TAD boundary stability across species to find human-specific structures. Finally, as high-resolution Hi-C becomes more prevalent from diverse tissues and individuals, we anticipate that computational prediction of personalized cell-type-specific TAD structure will facilitate understanding of how any possible variant is likely to affect 3D genome structure, gene regulation, and disease risk.

## METHODS

### Defining TADs

TAD maps for 37 different cell types were obtained from the 3D genome browser in BED format (Table S1) [44]. All TAD map analyses were conducted using the hg19 genome build. All cell types were available in hg19 format, except the Liver data, which we downloaded in hg38 and used the UCSC liftOver tool to convert to hg19 [66,67]. All TAD maps were systematically predicted from Hi-C data using the hidden Markov model (HMM) pipeline from Dixon *et al*. (2012) [19,44]. All analyses on the TAD map BED files were performed using the pybedtools wrapper for BedTools [68,69]. Most are maps defined with respect to the same 40 kb windows, except seven cell types (GM12878, HMEC, HUVEC, IMR90, K562, KBM7, NHEK) were defined using 25 kb windows.

### Quantifying partitioned heritability with S-LDSC

We conducted partitioned heritability using stratified-LD Score Regression v1.0.1 (S-LDSC) to test whether an annotation of interest (e.g. TADs or TAD boundaries) is enriched for heritability of a trait [43,49]. We considered GWAS summary statistics from a previously-described representative set of 41 diseases and complex traits [50,51,60,52–59]. Previous studies using these traits had GWAS replicates (genetic correlation > 0.9) for six of these traits. For these six traits (BMI, Height, High Cholesterol, Type 2 Diabetes, Smoking status, Years of Education), we considered only the GWAS with the largest sample size so our combined analysis did not overrepresent these six. All GWAS are European-ancestry only. We use 1000 Genomes for the LD reference panel (SNPs with MAF > 0.05 in European samples) [70] and HapMap Project Phase 3 (HapMap 3) [71] excluding the MHC region for our regression SNPs to estimate heritability enrichment and standardized effect size metrics.

#### Heritability enrichment

S-LDSC estimates the heritability enrichment, defined as the proportion of heritability explained by SNPs in the annotation divided by the proportion of SNPs in the annotation. The enrichment of annotation *c* is estimated as

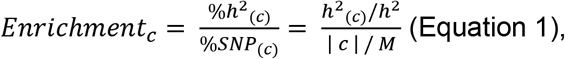

where *h^2^_(c)_* is the heritability explained by common SNPs in annotation *c*, *h^2^* is the heritability explained by the common SNPs over the whole genome, |*c* | is the number of common SNPs that lie in the annotation, and *M* is the number of common SNPs considered over the genome [43,51]. To investigate trends across all traits, we average the heritability enrichment and provide a confidence interval. When compared to meta-analysis using a random effects model conducted using Rmeta (function meta.summaries() [43,51,61,72] the trends are consistent (Fig. S1); therefore, we use averaging for future analyses to improve interpretability and reduce over-representation of higher-powered GWAS traits.

#### Standardized effect size

In contrast to heritability enrichment, standardized effect size (τ*_c_) quantifies effects that are unique to the focal annotation compared to a set of other annotations. The standardized effect size for annotation *c is*

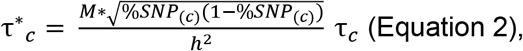

where τ_c_, estimated by LDSC, is the per-SNP contribution of (a one standard deviation increase in) annotation *c* to heritability, jointly modeled with other annotations.[51,62] The estimate of τc is conditioned on 86 diverse annotations from the baseline v2.1 model including coding, UTR, promoter and intronic regions, histone marks (H3K4me1, H3K4me3, H3K9ac, H3K27ac), DNAse I hypersensitivity sites (DHSs), chromHMM and Segway predictions, super-enhancers, FANTOM5 enhancers, GERP annotations, MAF bins, LD-related, and conservation annotations [43,62,63]. Standard errors are computed by LDSC using a block-jackknife. To combine across traits, we meta-analyze using a random effects model using Rmeta [72].

### Heritability enrichment across the TAD landscape

We analyzed TADs plus 50% of their total length on each side and subdivided into 20 equal-sized partitions, motivated by Krefting *et al* [25]. Hence, the center 10 bins (6-15) are inside the TAD. Bins 1-5 are upstream of the TAD and 16-20 are downstream of the TAD. We ran S-LDSR on these 20 bins across TAD maps from 37 cell types to calculate heritability enrichment over 41 traits. We investigated the heritability enrichment (or depletion) trends averaged across all traits and cell types (Fig. 2A), by cell type (Fig. 2B), and by trait (Fig. 4A). For the analyses by cell type and by trait, we clustered the heritability landscapes to determine if related cell types or related traits had similar patterns of heritability across the 3D genome. To do so, correlation distance was used as the distance metric and the clustering metric used average linkage. When clustering traits by their heritability landscape across the 3D genome, two agglomerative clusters were defined and termed “boundary-enriched” and “boundary-depleted”.

#### Using other TAD callers

To assess the influence of technical variation of TAD calling on these findings, we assessed the heritability patterns in human embryonic stem cells across TADs called by seven diverse methods (Armatus, Arrowhead, DomainCaller, HiCseg, TADbit, TADtree, TopDom). The TADs were called and published by Dali et al. 2017 [73] using Hi-C from Dixon et al. 2015 [35]. All other analysis was conducted the same as described using the 3DGenomeBrowser TADs.

### Sequence-level conservation across the TAD landscape

Using the same 20 partitions described above, we considered PhastCons element overlap to quantify evolutionary constraint across the TAD landscape. PhastCons elements are determined through a maximum likelihood fitting of a phylo-HMM across a group of 46 vertebrate genomes to predict conserved elements [74]. The BED file for these conserved elements was downloaded from the UCSC table browser in hg19 [67,75]. Each element has a score describing its level of conservation (a transformed log-odds score between 0 and 1000). This PhastCons element file was then intersected with bed files of the 20 partitions across the TAD landscape. Across each partition, we quantified the number of PhastCons base pairs per partition (regardless of score) and the average PhastCons element score per boundary (Fig. 2C).

### Quantifying boundary overlap and stability

For each cell type, we defined a set of boundaries with regards to the same windows across the genome. We considered boundaries defined by three different strategies.

#### 100 kb boundaries

We defined 100 kb boundaries (results shown in main text) to be regions 100 kb upstream of the TAD start and 100 kb downstream of the TAD end. For example, if a TAD was at chr1: 2,000,000-3,000,000, we would define its TAD boundaries to be at chr1:1,900,000-2,000,000 (boundary around the start) and chr1:3,000,000-3,100,000 (boundary around the end). Therefore, to quantify stability, we examined each 100 kb window across the genome. We removed boundaries that had any overlap with genomic gaps (centromeric/telomeric repeats from UCSC table browser) [67,75]. If there was a TAD boundary at that window for any of the cell types, we counted how many cell types (out of 37) shared that boundary. If only one cell type had a boundary at that location, it would be considered a “unique” boundary; whereas if it was observed in many cell types, it would be considered “stable (Fig. 3).

These boundaries were divided into quartiles of cell-type-stability. The cell type stability for quartile 1 included only boundaries observed in a single cell type, quartile 2 included boundaries in [2,4] cell types, quartile 3 included boundaries in [5,13], and quartile 4 included boundaries in 14 or more cell types (Fig. 3B).

#### 40 kb boundaries

We defined 40 kb boundaries (results in supplement) to be 40 kb windows surrounding (± 20 kb) TAD start and stop sites. For example, if a TAD was located at chr1:2,000,000-3,000,000, we would define its TAD boundaries to be at chr1:1,980,000-2,020,000 (boundary around the start) and chr1:2,980,000-3,020,000 (boundary around the end). We removed boundaries that had any overlap with genomic gaps (centromeric/telomeric repeats from UCSC table browser) [67,75]. These boundaries were divided into quartiles of cell-type-stability (Fig. S3A).

#### 200 kb boundaries

We defined 200 kb boundaries (results in supplement) to be 200 kb upstream of the TAD start and 200 kb downstream of the TAD end. For example, if a TAD was at chr1:2,000,000-3,000,000, we would define its TAD boundaries to be at chr1:1,800,000-2,000,000 (boundary around the start) and chr1:3,000,000-3,200,000 (boundary around the end). We removed boundaries that had any overlap with genomic gaps (centromeric/telomeric repeats from UCSC table browser) [67,75]. These boundaries were divided into quartiles of cell-type-stability (Fig. S3B).

### Quantifying TAD boundary profile similarity

To quantify TAD boundary profile similarity between any two cell types, we calculate a Jaccard similarity coefficient. We do this by counting the number of shared boundaries (intersection) and dividing by the total boundaries over both tissues (union). For the TAD boundary similarity heatmaps in Fig. S4, we clustered the cell types using complete-linkage (*i.e*. farthest neighbor) with the Jaccard distance metric (1-stability). The plots were made using Seaborn’s clustermap [76].

### Heritability enrichment by TAD boundary stability

S-LDSC was conducted on each quartile of stability for all 41 traits. Partitions for each quartile include TAD boundaries of that stability (see above). Enrichment values were log-scaled and linear regression (log10(heritability enrichment) ~ quartile of stability) was conducted. These results are shown in Figs. 3C, S5A, and S5D for 100 kb, 40 kb, and 200 kb boundary definitions respectively.

### TAD boundary stability and enrichment analyses

#### Evolutionary constraint

Evolutionary constraint was quantified by PhastCons elements, described above [74]. The PhastCons elements were intersected with the TAD boundary bed files, partitioned by stability. The two metrics of overlap quantification are the number of PhastCons base pairs per boundary regardless of score (base pairs per boundary) and the average PhastCons element score per boundary (average score of elements in the boundary).

#### CTCF enrichment

CTCF binding sites were determined through ChIP-seq analyses from ENCODE.[47,48] We searched for all CTCF ChIP-seq data with the following criteria: experiment, released, ChIP-seq, human (hg19), all tissues, adult, BED NarrowPeak file format, exclude any experiments with biosample treatments. Across all files, the CTCF peaks were concatenated, sorted, and merged into a single bed file. Overlapping peaks were merged into a single larger peak. The CTCF bed file was intersected with the TAD boundary bed files to quantify CTCF ChIP-seq peak’s overlap with each TAD boundary, stratified by that boundary’s stability. The two metrics of quantification were the number of CTCF ChIP-seq peaks per boundary (peaks per boundary) and the number of CTCF peak base pairs overlapping each boundary (base pairs per boundary).

#### Genes & Protein-coding genes

A list of all RefSeq genes and hg19 coordinates were downloaded from the UCSC table browser.[67,75,77] These were converted into a simplified list which included coordinates of only one transcript per gene (the longest) and included only autosomal and sex chromosome genes. From the simplified list of RefSeq genes, a subset of protein-coding genes was also created (identified using RefSeq accession numbers starting with NM). The simplified RefSeq gene list contains 27,090 genes. The simplified protein-coding RefSeq gene list contains 19,225 genes. These two gene sets were intersected with the TAD boundary bed files stratified by boundary stability. The metrics of overlap quantification are the number of genes or protein-coding genes per boundary.

#### Housekeeping genes

3804 housekeeping genes were downloaded from Eisenberg & Levanon (2013).[78] We converted the gene names into hg19 coordinates using RefSeq genes from the UCSC table browser, resulting in a BED file list of coordinates for 3681 genes (coordinates for a small number of genes were not found in the RefSeq list).[67,75,77] For each gene, coordinates of the longest transcript were considered. These housekeeping genes were intersected with the TAD boundary bed files stratified by boundary stability. The metrics of overlap quantification is the number of housekeeping genes or protein-coding genes per boundary.

### Defining GWAS phenotypic classes

To determine if similar phenotypes had similar heritability patterns across the 3D genome, we defined eight different phenotypic classes (Table S2): cardiopulmonary (N = 4), dermatologic (N = 7), hematologic (N = 5), immunologic (N = 4), metabolic (N = 7), neuropsychiatric (N = 8), reproductive (N = 4), and skeletal (N = 2). Our clusters originated from GWAS Atlas’ domains[79]; however, the categories were modified to place more emphasis on disease pathophysiology instead of involved organ system (*e.g*. Crohn’s Disease and Rheumatoid Arthritis were moved from the gastrointestinal and connective tissue categories respectively to an immunologic category). Similar categories were also combined (*e.g*. metabolic and endocrine, cardiovascular and respiratory).

### Data analysis and figure generation

Data and statistical analyses were conducted in Python 3.5.4 (Anaconda distribution) and R 3.6.1. Figure generation was aided by Matplotlib, Seaborn, and Inkscape.[76,80,81] This work was conducted in part using the resources of the Advanced Computing Center for Research and Education (ACCRE) at Vanderbilt University, Nashville, TN.

### Data Availability Statement

The publicly available datasets and software used for analysis are available in the following repositories: TAD maps from 3D Genome Browser [https://promoter.bx.psu.edu/hi-c/publications.html][44], LDSC [https://github.com/bulik/ldsc][43,49], GWAS traits formatted for LDSC from the Alkes Price lab [https://data.broadinstitute.org/alkesgroup/LDSCORE/independent_sumstats/], housekeeping genes [https://www.tau.ac.il/~elieis/HKG/HK_genes.txt][78], PhastCons elements, RefSeq Genes, and genome gaps [https://genome.ucsc.edu/cgi-bin/hgTables][67,74,75,77], and CTCF ChIP-seq peaks [https://www.encodeproject.org/][47,48].

The datasets we generated are available in the TAD-stability-heritability GitHub repository [https://github.com/emcarthur/TAD-stability-heritability] with DOI available at Zenodo[82] and include all results of our boundary calling (40 kb, 100 kb bookend, and 200 kb collapsed) and all partitioned heritability analysis output (by cell type and trait). Code is available upon request.

## Authors’ contributions

EM and JAC conceived and designed all the work presented here. EM conducted all the analysis. EM and JAC interpreted the results, drafted the work, and substantively revised the manuscript. EM and JAC have approved the submitted version and have agreed to be accountable for their contributions.

## Acknowledgements

The authors would like to thank Katherine S. Pollard, Emily Hodges, Geoff Fudenberg, Sarah Fong, Mary Lauren Benton, and other members of the Capra Lab for helpful discussions and manuscript comments. They would like to thank Margaux L.A. Hujoel and Steven Gazal for their help with LDSC implementation and heritability result interpretation. This work was conducted in part using the resources of the Advanced Computing Center for Research and Education at Vanderbilt University, Nashville, TN.

