## Supplemental Text, Figures S1-12, Tables S1-2 for "Topologically associating domain (TAD) boundaries stable across diverse cell types are evolutionarily constrained and enriched for heritability"

*TAD length*

The median TAD length across all cell types is 1.15 Mb (IQR: 0.71 - 1.82 Mb) and the median number of TADs per cell type is 1844 (IQR: 1625 - 2277). We observed an inverse relationship between TAD length and number of TADs in a cell type: cells with longer TADs have fewer TADs (Fig. S12). Primary tissues have longer TADs, whereas naïve cell types like stem cells and de-differentiated leukemia cell-lines have shorter TADs (Fig. S12). This is consistent with previous examination of neuronal development which found that, during differentiation, TAD number decreases with a corresponding increase in size [12].

*Similarity between TAD maps*

Our finding of TAD map similarity among functionally similar cell types contrasts with previous work by Sauerwald *et al.* (2018) which found that most similar TAD map pairs have no biological connection; however, they investigate a different set of cells (predominantly cancer cell lines) [13]. Comparisons with highly mutated cancer cell lines that may not reflect natural boundary patterns. Both our results and the Sauerwald comparisons could be influenced by batch effects because the Hi-C data considered were generated by different groups.

Nonetheless, our similarity quantifications agree with previous estimates. We find that the median pairwise Jaccard similarity for all 37 x 37 cell type comparisons is 0.18 (IQR: 0.15 - 0.23), 0.32 (IQR: 0.26 - 0.37), 0.41 (IQR: 0.35 - 0.47) at 40 kb, 100 kb, and 200 kb resolution, respectively. Our pairwise Jaccard similiarity between 200 kb boundaries (0.41) aligns with previous analyses that examined cell type TAD map similarity among larger windows have reported similarity coefficients between 0.4 - 0.5 [13]. At a finer resolution, Rao et al. (2014) reported Jaccard indices from 0.21 - 0.30 for comparisons of GM12878 to each of IMR90, HMEC, HUVEC, K562, KBM7 and NHEK [13,36]. The Jaccard similarity for our comparisons of these cell types is 0.24 - 0.37 (40 kb resolution).

Overall, this variability in TAD similarity across different cell types highlights the sensitivity of stability comparisons across to the definition of TAD boundaries used. For example, the median pairwise Jaccard similarity between 40 kb boundaries across 21 tissues defined by Schmitt et al. (2016) is 0.106 (IQR: 0.086 - 0.123). However, they collapsed boundaries to 200 kb “boundary regions” to conclude that TAD boundaries are highly stable (stating that over 35% of TAD boundaries are present in 21 of 21 tissues).[37] These previous studies often investigated more homogenous groups of cell types which could lead to higher estimates of stability. Ultimately, we stress than when interpreting claims of similarity between TAD maps of different cell types, the method of defining TADs (versus loop domains or boundary “regions”) and the breadth of cell types considered should be considered for context.

### SUPPLEMENTAL FIGURES

**Figure S1.** For TADs across 37 cell types, heritability is enriched near TAD boundaries when meta-analyzed across 41 common complex phenotypes. When combining data across traits, The heritability enrichment results are consistent using random-effects meta-analysis model (here) versus averaging ($R^2$ = 0.85, $p$ = 7x$10^{-9}$,Fig. 2A). The error band signifies a 99% confidence interval.

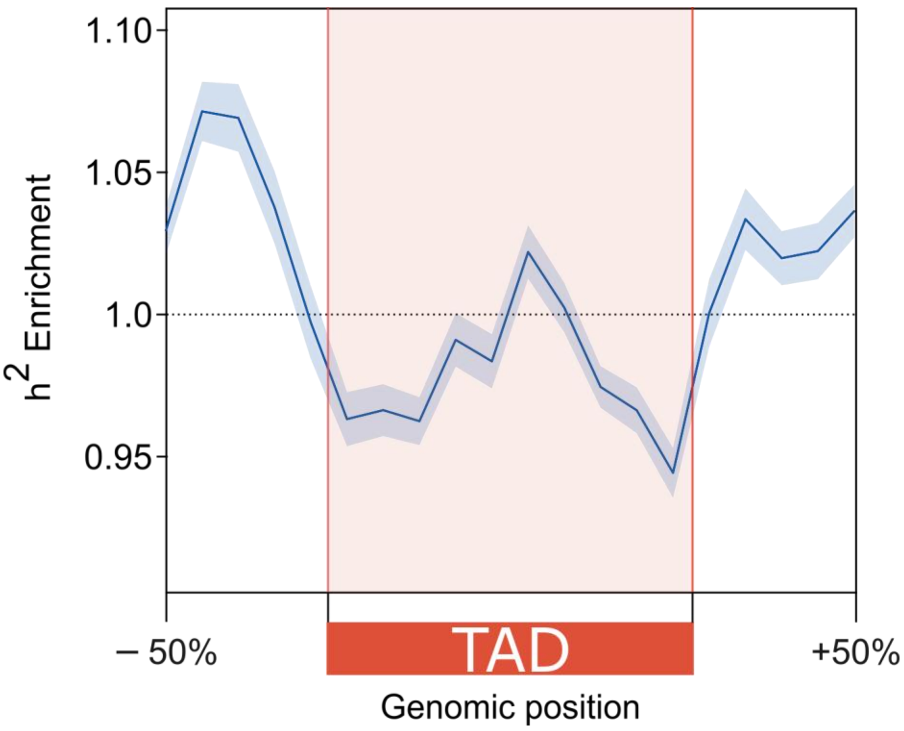

**Figure S2.** When meta-analyzed across all traits, there is no single part of the TAD landscape that is uniquely informative for trait heritability across all cell types when conditioned on a broad set of 86 functional (*e.g.* regulatory, conservation, coding, LD-related) annotations. Each line represents the standardized effect size meta-analyzed across all traits for that cell type (n = 37). The error bands signify 99% confidence intervals.

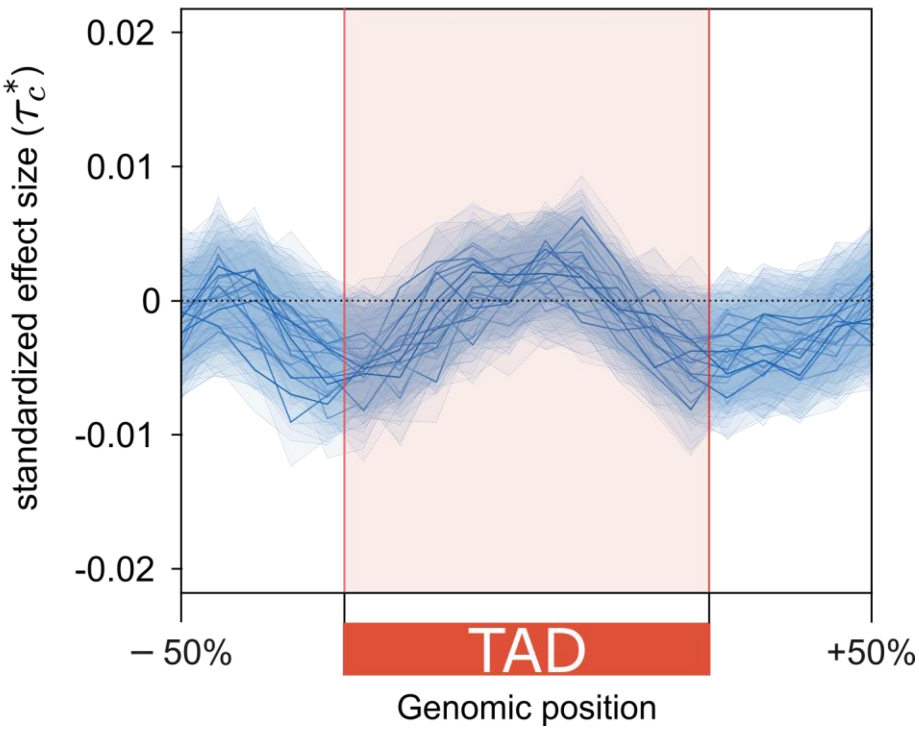

**Figure S3. Alternate definitions of TAD boundaries**. We show histograms of TAD boundaries by the number of cell types they are observed in (their "stability") colored by quartiles. In addition to the 100 kb boundary definitions (Fig. 3B), our supplemental analysis investigates **(A)** 40 kb boundaries and **(B)** 200 kb boundaries. Using the 40 kb definition, 33.9% of boundaries are unique to a single context and 2.0% of boundaries are observed in 25+ of 37 cell types. Using the 200 kb definition, 14.0% of boundaries are unique to a single context and 18.3% of boundaries are observed in 25+ of 37 cell types.

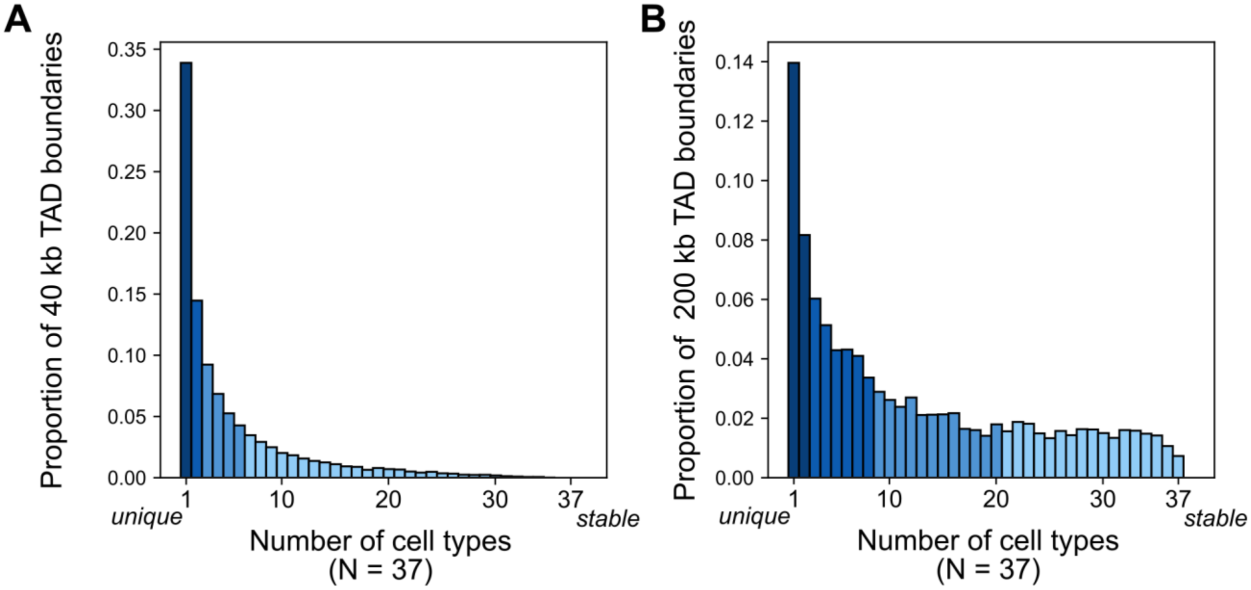

**Figure S4. Biologically similar cell types cluster by TAD map similarity**. Clustering for 37 cell types using the pairwise Jaccard similarity metric with colors labelling cellular classification groups for **(A)** 40 kb boundaries, **(B)** 100 kb boundaries, and **(C)** 200 kb boundaries.

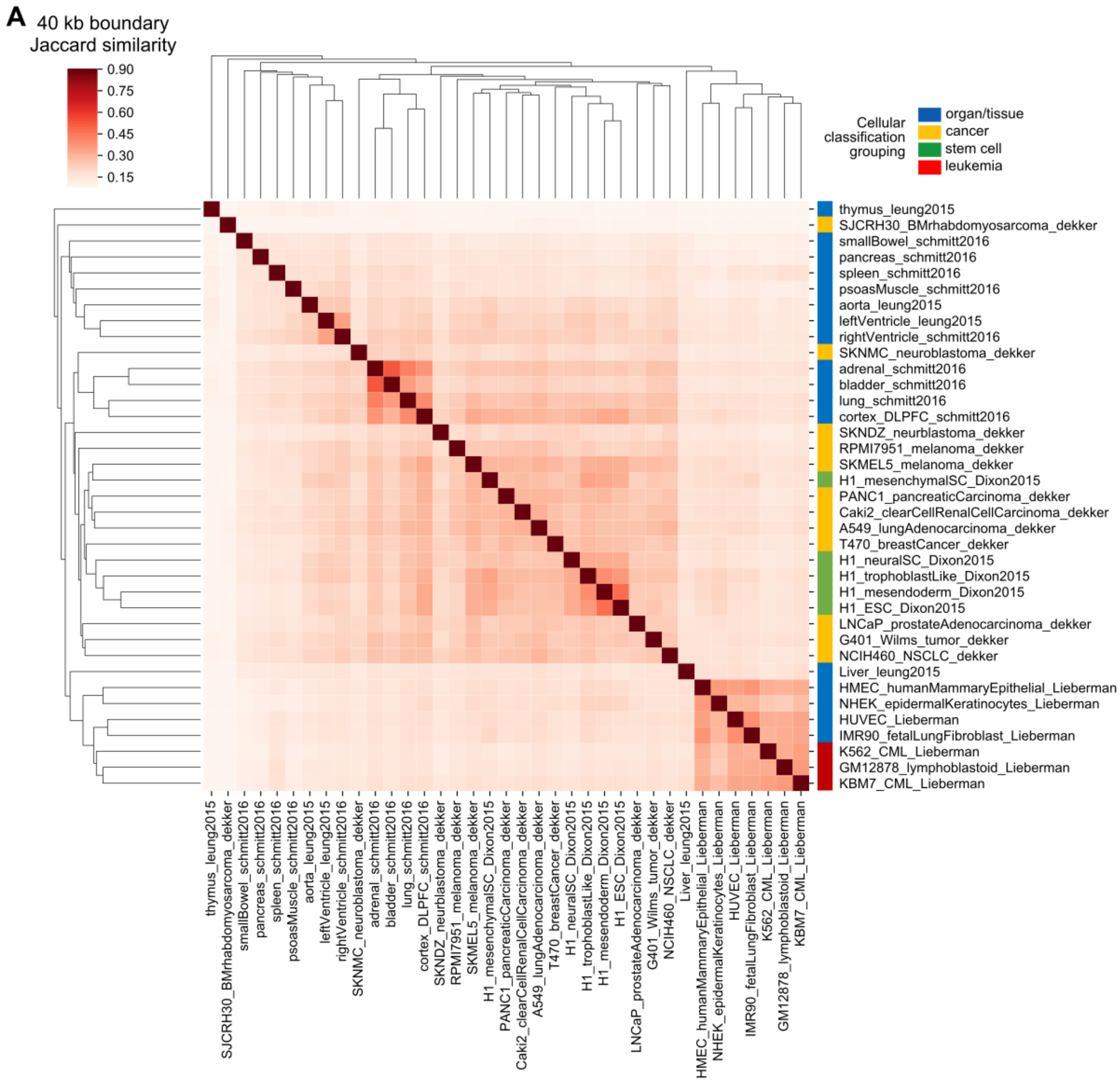

B

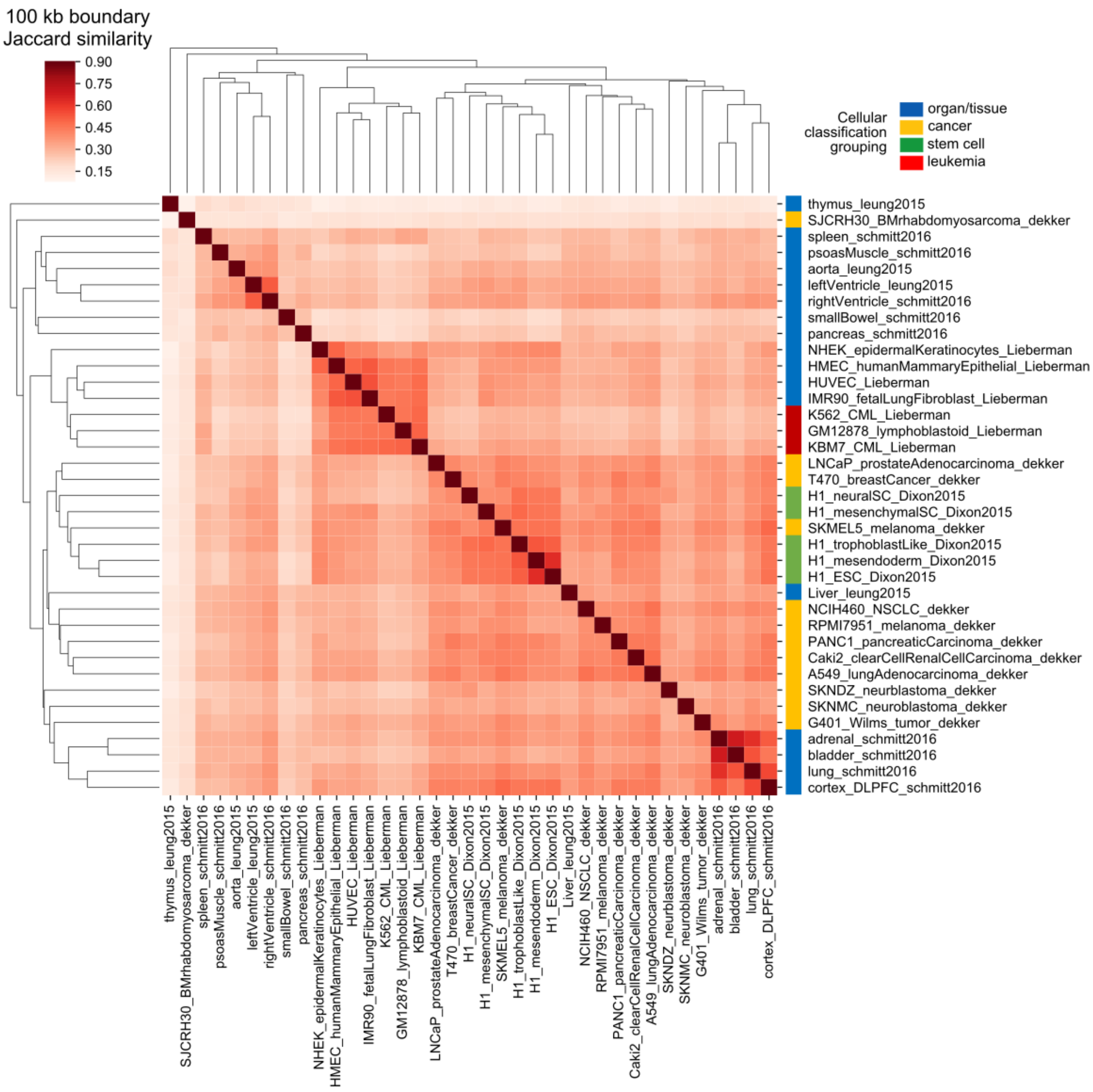

100 kb boundary
Jaccard similarity
0.90
0.75
0.60
0.45
0.30
0.15
Cellular
classification
grouping
organ/tissue
cancer
stem cell
leukemia
thymus_leung2015
SJCRH30_BMrhabdomyosarcoma_dekker
spleen_schmitt2016
psoasMuscle_schmitt2016
aorta_leung2015
leftVentricle_leung2015
rightVentricle_schmitt2016
smallBowel_schmitt2016
pancreas_schmitt2016
NHEK_epidermalKeratinocytes_Lieberman
HMEC_humanMammaryEpithelial_Lieberman
HUVEC_Lieberman
IMR90_fetalLungFibroblast_Lieberman
K562_CML_Lieberman
GM12878_lymphoblastoid_Lieberman
KBM7_CML_Lieberman
LNCaP_prostateAdenocarcinoma_dekker
T470_breastCancer_dekker
H1_neuralSC_Dixon2015
H1_mesenchymalSC_Dixon2015
SKMEL5_melanoma_dekker
H1_trophoblastLike_Dixon2015
H1_mesendoderm_Dixon2015
H1_ESC_Dixon2015
Liver_leung2015
NCIH460_NSCLC_dekker
RPMI7951_melanoma_dekker
PANC1_pancreaticCarcinoma_dekker
Caki2_clearCellRenalCellCarcinoma_dekker
A549_lungAdenocarcinoma_dekker
SKNDZ_neurblastoma_dekker
SKNMC_neuroblastoma_dekker
G401_Wilms_tumor_dekker
adrenal_schmitt2016
bladder_schmitt2016
lung_schmitt2016
cortex_DLPFC_schmitt2016

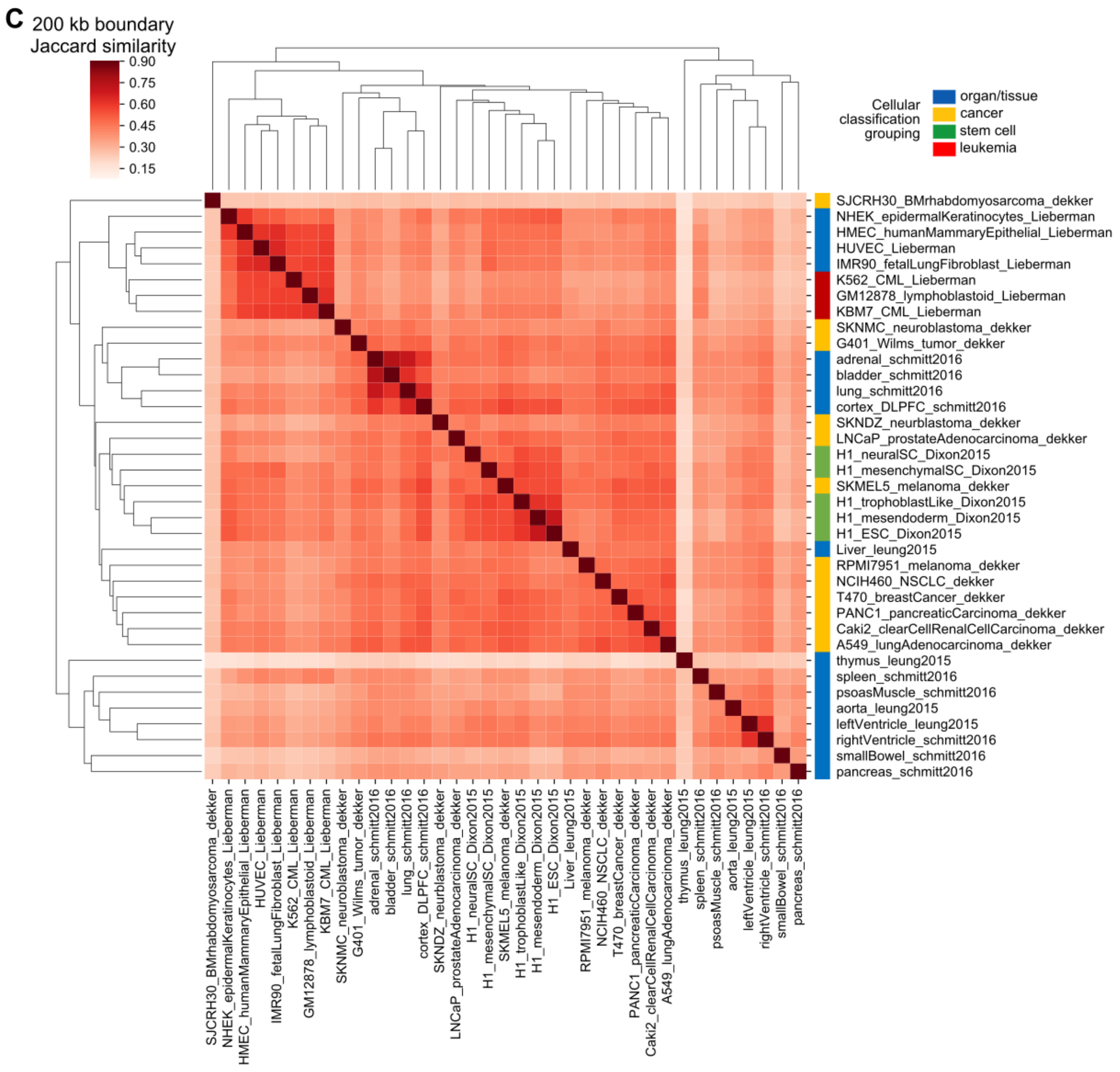
C
200 kb boundary
Jaccard similarity
0.90
0.75
0.60
0.45
0.30
0.15
Cellular classification grouping
organ/tissue
cancer
stem cell
leukemia
SJCRH30_BMrhabdomyosarcoma_dekker
NHEK_epidermalKeratinocytes_Lieberman
HMEC_humanMammaryEpithelial_Lieberman
HUVEC_Lieberman
IMR90_fetalLungFibroblast_Lieberman
K562_CML_Lieberman
GM12878_lymphoblastoid_Lieberman
KBM7_CML_Lieberman
SKNMC_neuroblastoma_dekker
G401_Wilms_tumor_dekker
adrenal_schmitt2016
bladder_schmitt2016
lung_schmitt2016
cortex_DLPFC_schmitt2016
SKNDZ_neurblastoma_dekker
LNCaP_prostateAdenocarcinoma_dekker
H1_neuralSC_Dixon2015
H1_mesenchymalSC_Dixon2015
SKMEL5_melanoma_dekker
H1_trophoblastLike_Dixon2015
H1_mesendoderm_Dixon2015
H1_ESC_Dixon2015
Liver_leung2015
RPMI7951_melanoma_dekker
NCIH460_NSCLC_dekker
T470_breastCancer_dekker
PANC1_pancreaticCarcinoma_dekker
Caki2_clearCellRenalCellCarcinoma_dekker
A549_lungAdenocarcinoma_dekker
thymus_leung2015
spleen_schmitt2016
psoasMuscle_schmitt2016
aorta_leung2015
leftVentricle_leung2015
rightVentricle_schmitt2016
smallBowel_schmitt2016
pancreas_schmitt2016

**Figure S5. Relationship between heritability enrichment and boundary stability is maintained despite different boundary definitions.** Over all traits, there is a positive relationship between boundary stability and heritability enrichment using 40 kb boundaries (**A**, *P* = 0.61), 100 kb boundaries (Fig. 3C, *P* = 0.006), and 200 kb boundaries (**D**, *P* = 2x10$^{-5}$). For traits in the boundary-enriched cluster (Fig. 4B), there is a stronger positive relationship between boundary stability and heritability using 40 kb boundaries (**B**, *P* = 0.06), 100 kb boundaries (Fig. 4D, *P* = 2x10$^{-6}$), and 200 kb boundaries (**E**, *P* = 3x10$^{-14}$). For traits in the boundary-depleted cluster (Fig. 4C), there is a weak negative relationship between boundary stability and heritability using 40 kb boundaries (**C**, *P* = 0.09), 100 kb boundaries (Fig. 4F, *P* = 0.09), and 200 kb boundaries (**F**, *P* = 0.01). Error bars/bands signify 95% confidence intervals.

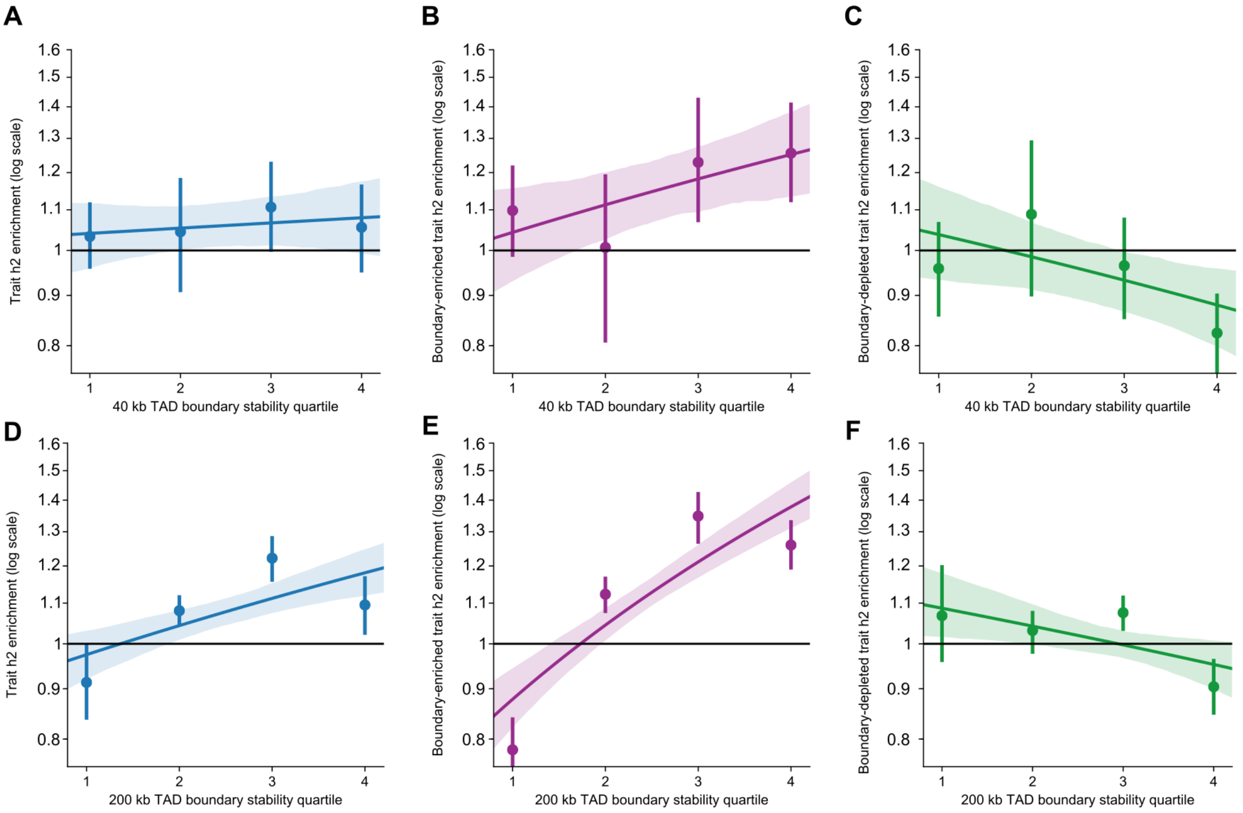

**Figure S6. Stable TAD boundaries are enriched for genes despite varying boundary definitions.** We show the relationship between increased TAD boundary stability and gene overlap using 40 kb boundaries **(A,D,G)**, 100 kb boundaries **(B,E,H)**, and 200 kb boundaries **(C,F,I)**. We also demonstrate this trend using three types of genes: all RefSeq genes **(A-C)**, protein-coding genes **(D-F)**, and housekeeping genes **(G-I)**. Panel H is shown in the main text (Fig. 3F). TAD boundary stability quartiles are defined by the empirical distributions shown in Fig. S3A (40 kb), Fig. 3B (100 kb), and Fig. S3B. Boundaries in the first quartile are unique to a single cell type, while boundaries in higher quartiles are stable across multiple cell types. Error bars/bands signify 95% confidence intervals.

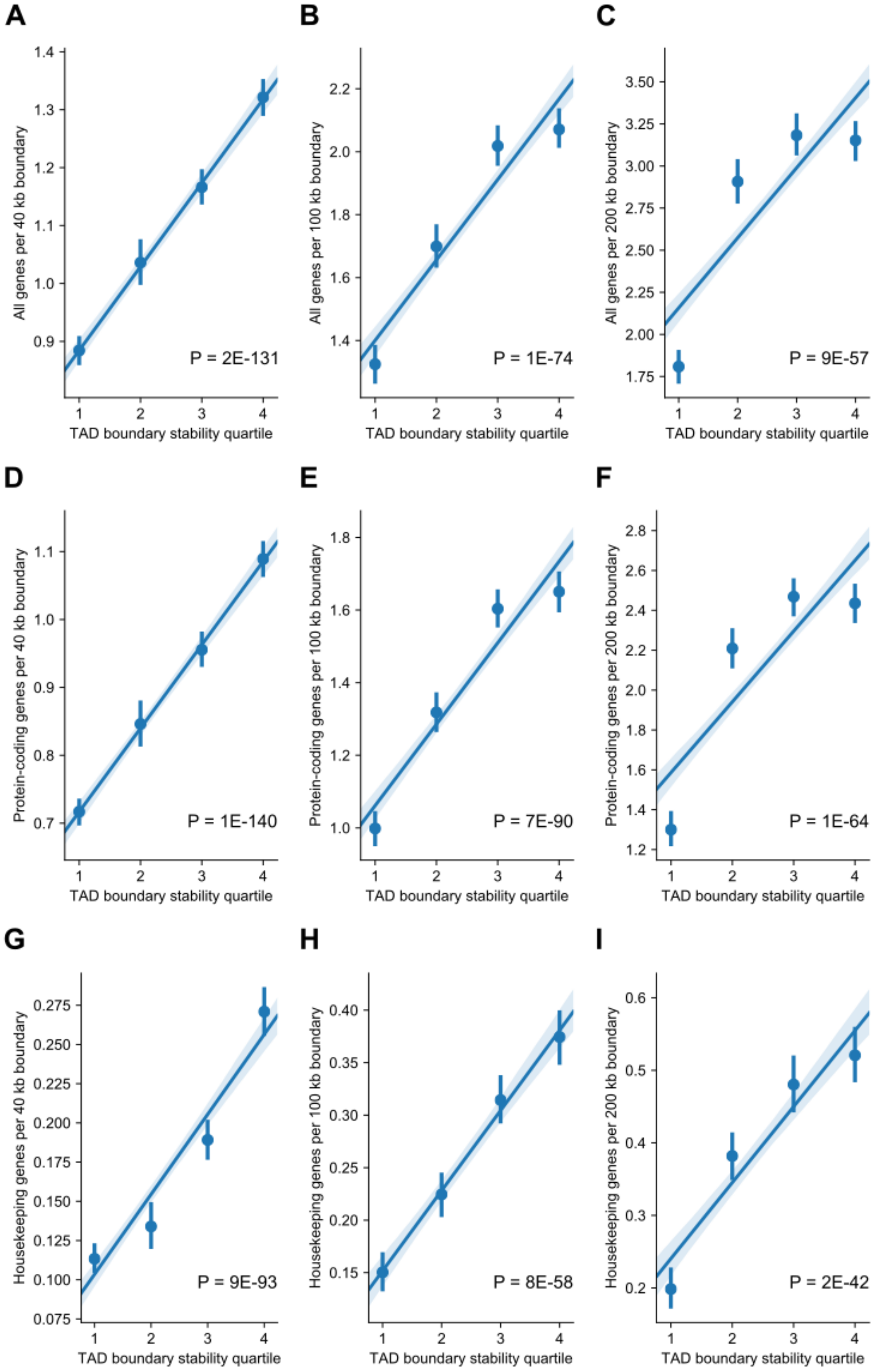

**Figure S7. Stable TAD boundaries are enriched for sequence-level conservation despite varying boundary definitions.** We show the relationship between increased TAD boundary stability and sequence-level conservation (via PhastCons element overlap) using 40 kb boundaries **(A & D)**, 100 kb boundaries **(B & E)**, and 200 kb boundaries **(C & F)**. We also demonstrate this trend holds at two different metrics of conservation: number of bases overlapping PhastCons elements **(A-C)** and average PhastCons element score per boundary **(D-F)**. Panel B is shown in the main text (Fig. 3D). TAD boundary stability quartiles are defined by the empirical distributions shown in Fig. S3A (40 kb), Fig. 3B (100 kb), and Fig. S3B. Boundaries in the first quartile are unique to a single cell type, while boundaries in higher quartiles are stable across multiple cell types. Error bars/bands signify 95% confidence intervals.

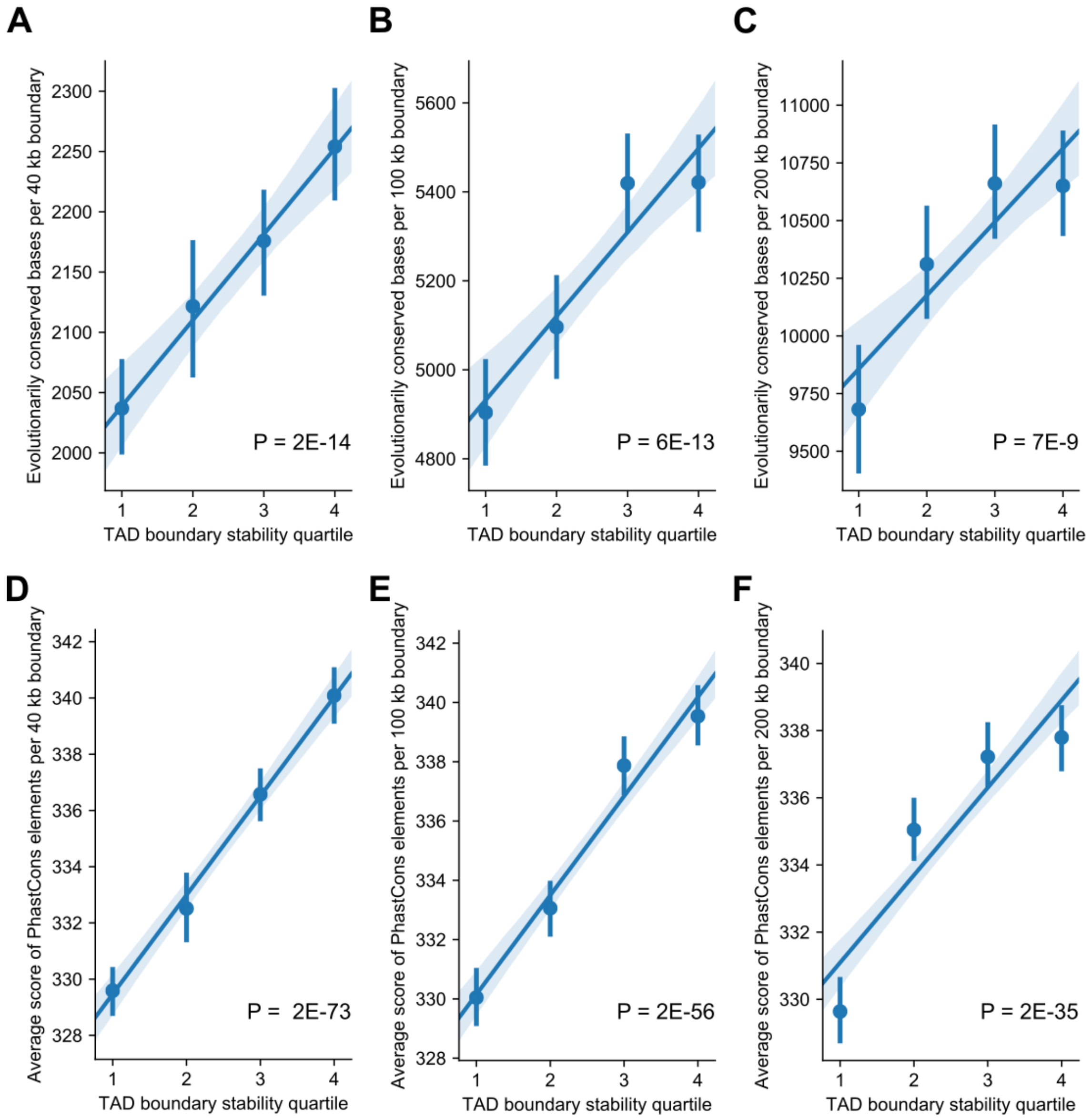

**Figure S8. Stable TAD boundaries are enriched for CTCF binding despite varying boundary definitions.** We show the relationship between increased TAD boundary stability and CTCF binding using 40 kb boundaries **(A & D)**, 100 kb boundaries **(B & E)**, and 200 kb boundaries **(C & F)**. We also demonstrate this trend holds at two different metrics of CTCF overlap: count of CTCF ChIP-seq peaks per boundary **(A-C)** and number of CTCF ChIP-seq peak bases overlapping each boundary **(D-F)**. Panel B is shown in the main text (Fig. 3E). TAD boundary stability quartiles are defined by the empirical distributions shown in Fig. S3A (40 kb), Fig. 3B (100 kb), and Fig. S3B. Boundaries in the first quartile are unique to a single cell type, while boundaries in higher quartiles are stable across multiple cell types. Error bars/bands signify 95% confidence intervals.

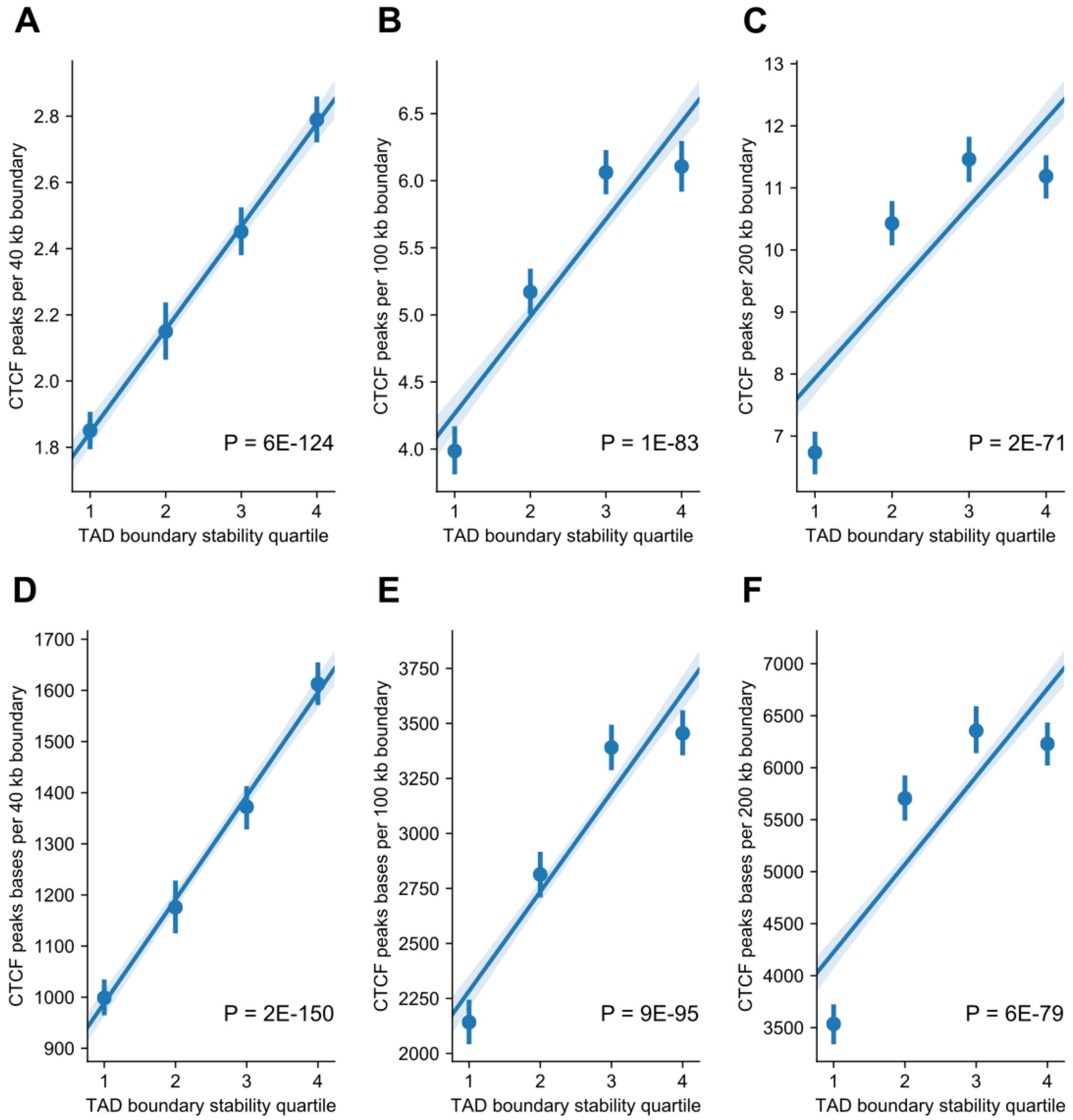

**Figure S9. Traits in the boundary-depleted cluster and boundary-enriched cluster do not differ for GWAS parameters including: (A)** Number of GWAS SNPs (p = 0.78, t-test with equal variances), **(B)** Number of individuals in the GWAS (p = 0.92), or **(C)** The traits SNP-based heritability (p = 0.88). Error bars signify 95% confidence intervals.

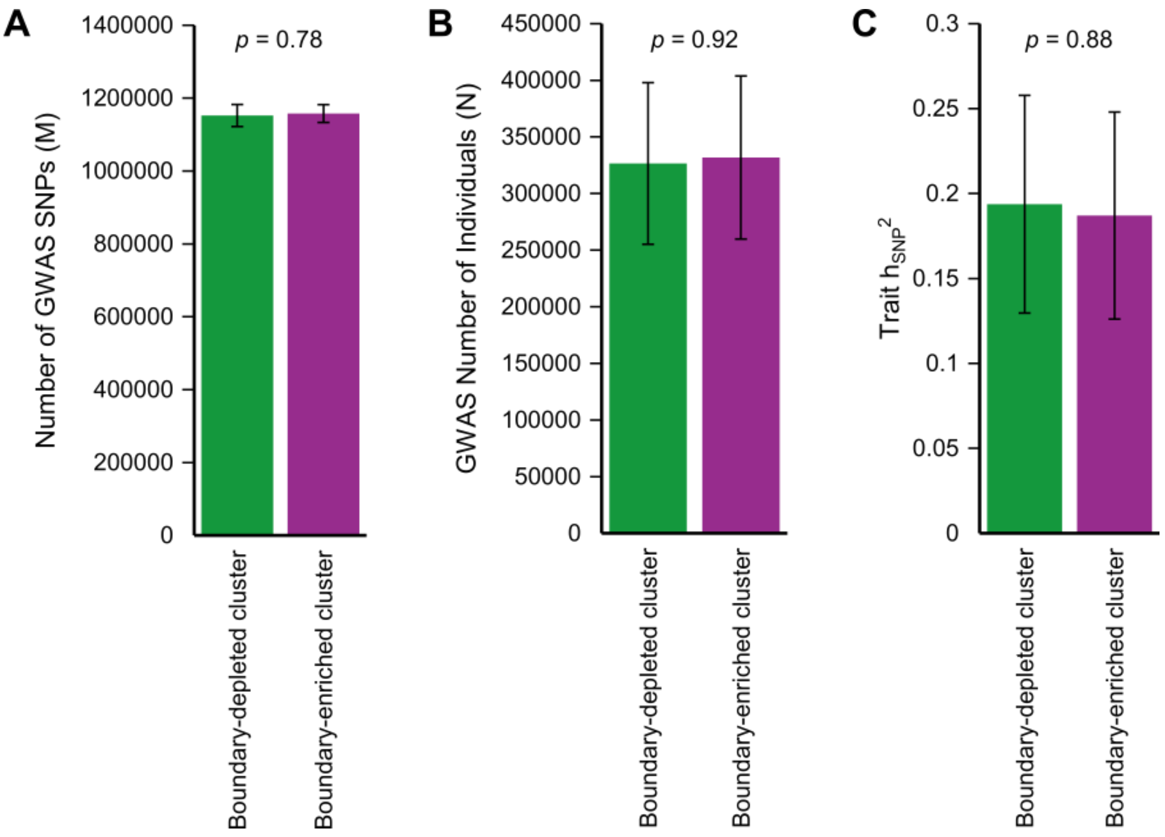

**Figure S10. Patterns of heritability enrichment across the 3D genome in human embryonic stem cells (ESC) are robust to the TAD definition algorithm used. (A)** Heritability enrichment landscape over TADs in ESCs called by eight different algorithms for traits in the boundary-enriched cluster. Like findings shown in Fig. 4B (which use TADs from the Dixon pipeline), TAD boundaries are enriched for heritability compared to TADs. **(B)** Heritability enrichment landscape over TADs in ESCs for traits in the boundary-depleted cluster. Like findings shown in Fig. 4C (which use TADs from the Dixon pipeline), TADs are centrally enriched for heritability compared to TAD boundaries. Error bands signify 95% confidence intervals.

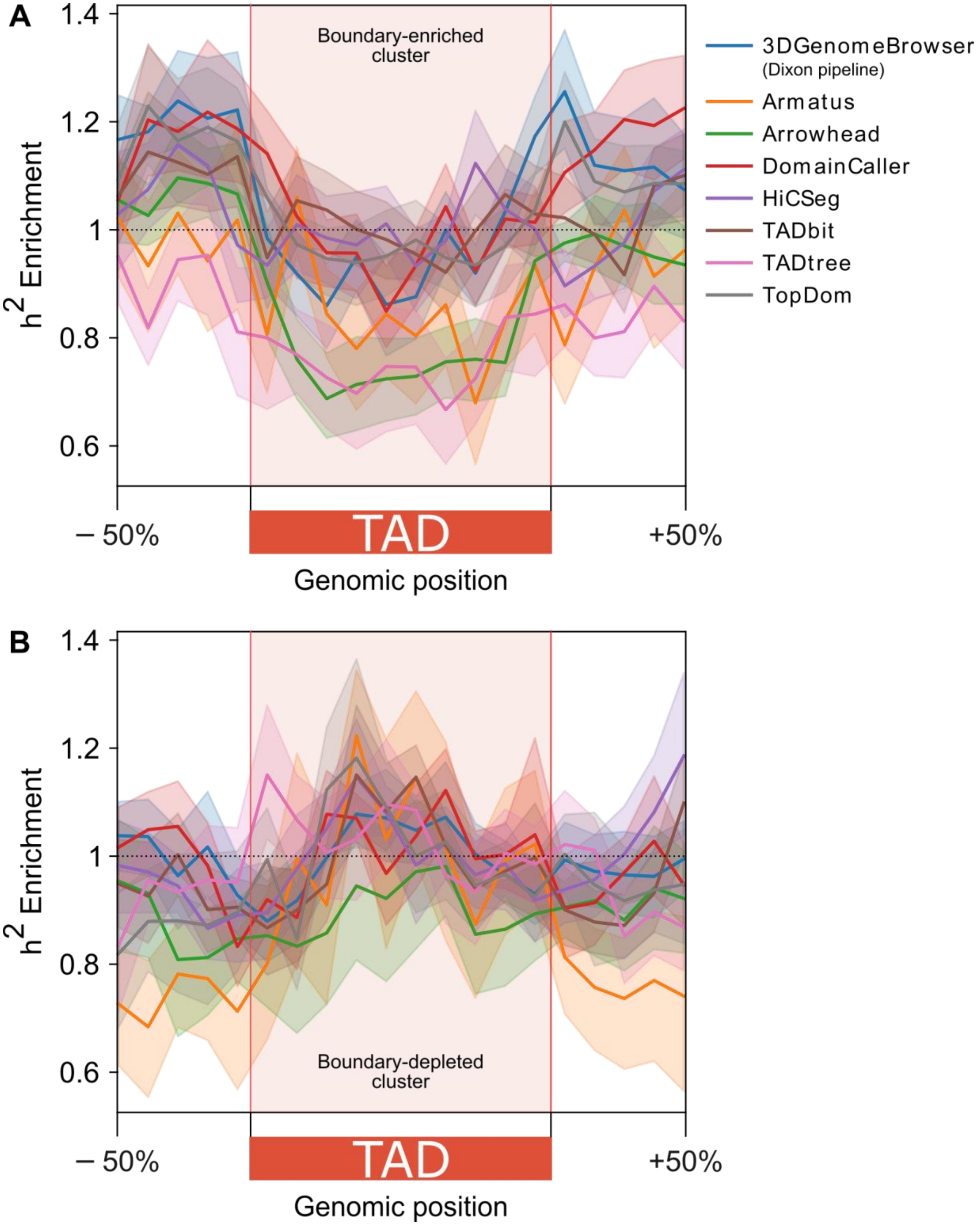

**Figure S11.** For the boundary-depleted cluster traits, TADs flanked by the most stable boundaries (measured by taking the average stability of its two boundaries and binning into quintiles) have increased heritability centrally. This analysis was performed in a random subset of 7 cell types (aorta, H1_ESC, leftVentricle, Liver, psoasMuscle, SKNDZ, T470). Error bands signify 95% confidence intervals.

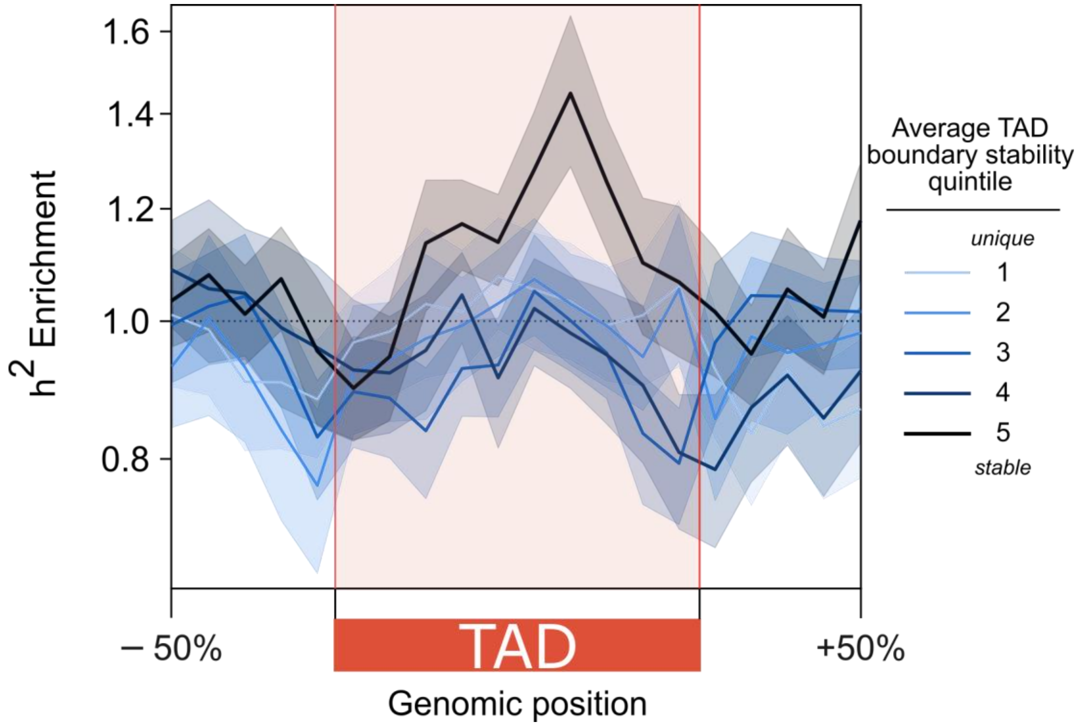

**Figure S12.** Across 37 cell types, there is an inverse relationship between TAD length and number of TADs. Organ/tissue cell types generally have the longest (and fewest) TADs. Leukemia and stem cells have the shortest (and most) TADs. Error bands signify the IQR.

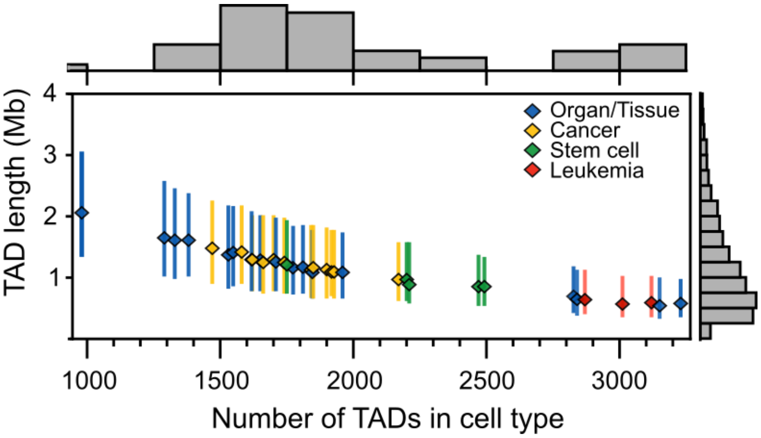

### Supplemental Table 1. Cell types used for all analyses from the 3DGenomeBrowser

| FileNameFrom3DGenomeBrowser | CellTypeDescription | Abbreviation | BiologicalCluster | Citation |
|---|---|---|---|---|
| A549_raw-rep1_TADs.txt | A549_lungAdenocarcinoma_dekker | A549 | cancer | Lajoie, Dekker et al. (2015)[46], ENCODE[47,48] |
| AdrenalGland_Donor-AD2-raw_TADs.txt | adrenal_schmitt2016 | adrenal | organ/tissue | Schmitt et al. (2016)[37] |
| Aorta_STL002_Leung2015-raw_TADs.txt | aorta_leung2015 | aorta | organ/tissue | Leung et al. (2015)[45] |
| Bladder_Donor-BL1-raw_TADs.txt | bladder_schmitt2016 | bladder | organ/tissue | Schmitt et al. (2016)[37] |
| Bowel_Small_Donor-SB2-raw_TADs.txt | smallBowel_schmitt2016 | smallBowel | organ/tissue | Schmitt et al. (2016)[37] |
| Caki2_raw-rep1_TADs.txt | Caki2_clearCellRenalCellCarcinoma_dekker | Caki2 | cancer | Lajoie, Dekker et al. (2015)[46], ENCODE[47,48] |
| Cortex_DLPFC_Donor-CO-raw_TADs.txt | cortex_DLPFC_schmitt2016 | DLPFC | organ/tissue | Schmitt et al. (2016)[37] |
| G401_raw-rep1_TADs.txt | G401_Wilms_tumor_dekker | G401 | cancer | Lajoie, Dekker et al. (2015)[46], ENCODE[47,48] |
| GM12878_Lieberman-raw_TADs.txt | GM12878_lymphoblastoid_Lieberman | GM12878 | leukemia | Rao et al. (2014)[36] |
| H1-ESC_Dixon2015-raw_TADs.txt | H1_ESC_Dixon2015 | ESC | stem cell | Dixon et al. (2015)[35] |
| H1-MES_Dixon2015-raw_TADs.txt | H1_mesendoderm_Dixon2015 | MES | stem cell | Dixon et al. (2015)[35] |
| H1-MSC_Dixon2015-raw_TADs.txt | H1_mesenchymalSC_Dixon2015 | MSC | stem cell | Dixon et al. (2015)[35] |
| H1-NPC_Dixon2015-raw_TADs.txt | H1_neuralSC_Dixon2015 | NPC | stem cell | Dixon et al. (2015)[35] |
| H1-TRO_Dixon2015-raw_TADs.txt | H1_trophoblastLike_Dixon2015 | TRO | stem cell | Dixon et al. (2015)[35] |
| HMEC_Lieberman-raw_TADs.txt | HMEC_humanMammaryEpithelial_Lieberman | HMEC | organ/tissue | Rao et al. (2014)[36] |
| HUVEC_Lieberman-raw_TADs.txt | HUVEC_Lieberman | HUVEC | organ/tissue | Rao et al. (2014)[36] |
| IMR90_Lieberman-raw_TADs.txt | IMR90_fetalLungFibroblast_Lieberman | IMR90 | organ/tissue | Rao et al. (2014)[36] |
| K562_Lieberman-raw_TADs.txt | K562_CML_Lieberman | K562 | leukemia | Rao et al. (2014)[36] |
| KBM7_Lieberman-raw_TADs.txt | KBM7_CML_Lieberman | KBM7 | leukemia | Rao et al. (2014)[36] |
| Liver_STL011_Leung_2015-raw_TADs_hg19From38.txt | Liver_leung2015 | Liver | organ/tissue | Leung et al. (2015)[45] |
| LNCaP_raw-rep1_TADs.txt | LNCaP_prostateAdenocarcinoma_dekker | LNCaP | cancer | Lajoie, Dekker et al. (2015)[46], ENCODE[47,48] |
| Lung_Donor-LG1-raw_TADs.txt | lung_schmitt2016 | lung | organ/tissue | Schmitt et al. (2016)[37] |
| Muscle_Psoas_Donor-PO1-raw_TADs.txt | psoasMuscle_schmitt2016 | psoas | organ/tissue | Schmitt et al. (2016)[37] |
| NCIH460_raw-rep1_TADs.txt | NCIH460_NSCLC_dekker | NCIH460 | cancer | Lajoie, Dekker et al. (2015)[46], ENCODE[47,48] |
| NHEK_Lieberman-raw_TADs.txt | NHEK_epidermalKeratinocytes_Lieberman | NHEK | organ/tissue | Rao et al. (2014)[36] |
| PANC1_raw-rep1_TADs.txt | PANC1_pancreaticCarcinoma_dekker | PANC1 | cancer | Lajoie, Dekker et al. (2015)[46], ENCODE[47,48] |
| Pancreas_Donor-PA2-raw_TADs.txt | pancreas_schmitt2016 | pancreas | organ/tissue | Schmitt et al. (2016)[37] |
| RPMI7951_raw-rep1_TADs.txt | RPMI7951_melanoma_dekker | RPMI7951 | cancer | Lajoie, Dekker et al. (2015)[46], ENCODE[47,48] |
| SJCRH30_raw-rep1_TADs.txt | SJCRH30_BMrhabdomyosarcoma_dekker | SJCRH30 | cancer | Lajoie, Dekker et al. (2015)[46], ENCODE[47,48] |
| SKMEL5_raw-rep1_TADs.txt | SKMEL5_melanoma_dekker | SKMEL5 | cancer | Lajoie, Dekker et al. (2015)[46], ENCODE[47,48] |
| SKNDZ_raw-rep1_TADs.txt | SKNDZ_neurblastoma_dekker | SKNDZ | cancer | Lajoie, Dekker et al. (2015)[46], ENCODE[47,48] |
| SKNMC_raw-rep1_TADs.txt | SKNMC_neuroblastoma_dekker | SKNMC | cancer | Lajoie, Dekker et al. (2015)[46], ENCODE[47,48] |
| Spleen_Donor-PX1-raw_TADs.txt | spleen_schmitt2016 | spleen | organ/tissue | Schmitt et al. (2016)[37] |
| T470_raw-rep1_TADs.txt | T470_breastCancer_dekker | T470 | cancer | Lajoie, Dekker et al. (2015)[46], ENCODE[47,48] |
| Thymus_STL001_Leung2015-raw_TADs.txt | thymus_leung2015 | thymus | organ/tissue | Leung et al. (2015)[45] |
| VentricleLeft_STL003_Leung2015-raw_TADs.txt | leftVentricle_leung2015 | leftVentricle | organ/tissue | Leung et al. (2015)[45] |
| VentricleRight_Donor-RV3-raw_TADs.txt | rightVentricle_schmitt2016 | rightVentricle | organ/tissue | Schmitt et al. (2016)[37] |

**Supplemental Table 2. Genome-wide association study (GWAS) traits used for heritability analyses**

| Nickname | Trait | M | h2 | h2_SE | N | Phenotypic class | actual cluster | Source |
|---|---|---|---|---|---|---|---|---|
| Anorexia | Anorexia | 931184 | 0.2153 | 0.0169 | 32143 | Neuropsych | Boundary-depleted | Boraska et al. 2014 Mol Psych[53] |
| ASD | Autism_Spectrum | 1173307 | 0.4607 | 0.0517 | 10263 | Neuropsych | Boundary-depleted | PGC Cross-Disorder Group, 2013 Lancet[54] |
| AutoimmuneDz | Auto_Immune_Traits_(Sure) | 1187056 | 0.0068 | 0.0013 | 459324 | Immunologic | Boundary-enriched | UKBiobank[52] |
| Balding | Balding_Type_I | 1187056 | 0.2154 | 0.019 | 208336 | Dermatologic | Boundary-depleted | UKBiobank[52] |
| BMI | BMI | 1187056 | 0.252 | 0.0071 | 457824 | Metabolic | Boundary-depleted | UKBiobank[52] |
| CrohnsDz | Crohn's_Disease | 1051514 | 0.4723 | 0.0575 | 20883 | Immunologic | Boundary-enriched | Jostins et al., 2012 Nature[55] |
| DepressiveSxs | Depressive_symptoms | 1115393 | 0.0473 | 0.0037 | 161460 | Neuropsych | Boundary-depleted | Okbay et al., 2016 Nat Genet[56] |
| DermDz | Dermatologic_Diseases | 1187056 | 0.0094 | 0.0014 | 459324 | Dermatologic | Boundary-enriched | UKBiobank[52] |
| Eczema | Eczema | 1187056 | 0.0675 | 0.0038 | 458699 | Dermatologic | Boundary-enriched | UKBiobank[52] |
| EosinophilCount | Eosinophil_Count | 1187056 | 0.1977 | 0.0143 | 439938 | Hematologic | Boundary-enriched | UKBiobank[52] |
| FEV1_FVC_Ratio | FEV1-FVC_Ratio | 1187056 | 0.2336 | 0.0113 | 371949 | Cardiopulmonary | Boundary-enriched | UKBiobank[52] |
| FirstBirthAge | Age_first_birth | 1079424 | 0.0617 | 0.0033 | 222037 | Reproductive | Boundary-depleted | Barban et al., 2016 Nat Genet[57] |
| FVC | Forced_Vital_Capacity_(FVC) | 1187056 | 0.2068 | 0.0065 | 371949 | Cardiopulmonary | Boundary-enriched | UKBiobank[52] |
| HairColor | Hair_Color | 1187056 | 0.4523 | 0.1497 | 452720 | Dermatologic | Boundary-depleted | UKBiobank[52] |
| HDL | HDL | 1019272 | 0.1362 | 0.0166 | 99900 | Metabolic | Boundary-enriched | Teslovich et al., 2010 Nature[58] |
| Heel_T_Score | Heel_T_Score | 1187056 | 0.3628 | 0.0307 | 445921 | Skeletal | Boundary-depleted | UKBiobank[52] |
| Height | Height | 1187056 | 0.6034 | 0.027 | 458303 | Skeletal | Boundary-enriched | UKBiobank[52] |
| HighCholesterol | High_Cholesterol | 1187056 | 0.0468 | 0.0039 | 459324 | Metabolic | Boundary-enriched | UKBiobank[52] |
| Hypothyroidism | Hypothyroidism | 1187056 | 0.0459 | 0.0037 | 459324 | Metabolic | Boundary-enriched | UKBiobank[52] |
| LDL | LDL | 1017973 | 0.121 | 0.0166 | 95454 | Metabolic | Boundary-enriched | Teslovich et al., 2010 Nature[58] |
| MenarcheAge | Age_at_Menarche | 1187056 | 0.2457 | 0.0102 | 242278 | Reproductive | Boundary-enriched | UKBiobank[52] |
| MenopauseAge | Age_at_Menopause | 1187056 | 0.1215 | 0.0086 | 143025 | Reproductive | Boundary-enriched | UKBiobank[52] |
| MorningPerson | Morning_Person | 1187056 | 0.1002 | 0.0035 | 410520 | Neuropsych | Boundary-depleted | UKBiobank[52] |
| Neuroticism | Neuroticism | 1187056 | 0.1113 | 0.0037 | 372066 | Neuropsych | Boundary-depleted | UKBiobank[52] |
| NumChildrenBorn | Number_children_ever_born | 1080059 | 0.0256 | 0.0018 | 318863 | Reproductive | Boundary-depleted | Barban et al., 2016 Nat Genet[57] |
| PlateletCount | Platelet_Count | 1187056 | 0.349 | 0.0294 | 444382 | Hematologic | Boundary-enriched | UKBiobank[52] |
| RA | Rheumatoid_Arthritis | 1125155 | 0.1694 | 0.023 | 38242 | Immunologic | Boundary-enriched | Okada et al., 2014 Nature[59] |
| RBCCount | Red_Blood_Cell_Count | 1187056 | 0.2434 | 0.0191 | 445174 | Hematologic | Boundary-enriched | UKBiobank[52] |
| RDW | Red_Blood_Cell_Distribution_Width | 1187056 | 0.2234 | 0.0198 | 442700 | Hematologic | Boundary-enriched | UKBiobank[52] |
| Resp_ENT_Dz | Respiratory_and_Ear-nose-throat_Diseases | 1187056 | 0.0483 | 0.0034 | 459324 | Cardiopulmonary | Boundary-depleted | UKBiobank[52] |
| Schizophrenia | Schizophrenia | 1083014 | 0.4512 | 0.0189 | 70100 | Neuropsych | Boundary-depleted | SCZ Working Group of the PGC, 2014 Nature[60] |
| SkinColor | Skin_Color | 1187056 | 0.1896 | 0.0539 | 453609 | Dermatologic | Boundary-depleted | UKBiobank[52] |
| SmokingStatus | Smoking_Status | 1187056 | 0.0972 | 0.0032 | 457683 | Neuropsych | Boundary-depleted | UKBiobank[52] |
| Sunburn | Sunburn_Occasion | 1187056 | 0.0915 | 0.0162 | 344229 | Dermatologic | Boundary-depleted | UKBiobank[52] |
| SystolicBP | Systolic_Blood_Pressure | 1187056 | 0.1966 | 0.007 | 422771 | Cardiopulmonary | Boundary-depleted | UKBiobank[52] |
| T2D | Type_2_Diabetes | 1187056 | 0.043 | 0.0025 | 459324 | Metabolic | Boundary-enriched | UKBiobank[52] |
| Tanning | Tanning | 1187056 | 0.172 | 0.0609 | 449984 | Dermatologic | Boundary-depleted | UKBiobank[52] |
| UC | Ulcerative_Colitis | 1076834 | 0.2424 | 0.032 | 27432 | Immunologic | Boundary-enriched | Jostins et al., 2012 Nature[55] |
| WaistHipRatio | Waist-hip_Ratio | 1187056 | 0.1423 | 0.0067 | 458417 | Metabolic | Boundary-enriched | UKBiobank[52] |
| WBCCount | White_Blood_Cell_Count | 1187056 | 0.1873 | 0.0105 | 444502 | Hematologic | Boundary-enriched | UKBiobank[52] |
| YearsOfEd | College_Education | 1187056 | 0.1299 | 0.0037 | 454813 | Neuropsych | Boundary-depleted | UKBiobank[52] |
